# CDH3 as a Novel Therapeutic Target in Basal-like Double-Negative Prostate Cancer

**DOI:** 10.1101/2025.11.18.689150

**Authors:** Guoqiang Liu, Shiyu Wang, Loan Duong, Ziyu Zeng, Jing Yang, Yun Zhao, Sharif Rahmy, Shan Feng, Ariana Arce, Linchuan Jia, Jun Wan, Liang Cheng, Xuemin Lu, Xin Lu

**Affiliations:** Integrated Biomedical Science Graduate Program, University of Notre Dame, Notre Dame, IN 46556, USA; Department of Biological Sciences, University of Notre Dame, Notre Dame, IN 46556, USA; iSURE Program, University of Notre Dame, Notre Dame, IN 46556, USA; Department of Medical and Molecular Genetics, Center for Computational Biology and Bioinformatics, Indiana University School of Medicine, Indianapolis, IN 46202, USA; Department of BioHealth Informatics, Luddy School of Informatics and Computing, Indiana University at Indianapolis, Indianapolis, IN 46202, USA; Indiana University Simon Comprehensive Cancer Center, Indiana University School of Medicine, Indianapolis, IN 46202, USA; Department of Pathology and Laboratory Medicine, Department of Surgery, Brown University Warren Alpert Medical School, the Legorreta Cancer Center at Brown University, and Brown University Health, Providence, RI, USA

## Abstract

**Purpose:** Basal-like (also known as double-negative) prostate cancers are aggressive tumors that lack effective targeted therapies. We aimed to delineate the role of CDH3 (P-cadherin) in basal-like prostate cancer and evaluate CDH3-directed therapeutic strategies.

**Methods:** We integrated genetically engineered mouse models (GEMMs) of prostate cancer, bulk and single-cell transcriptomic analyses, and a suite of in vitro and in vivo experiments. CDH3 expression and associated signaling pathways were examined in *Pten/Apc* double-knockout mouse prostates and human datasets. Functional studies included antibody-drug conjugate (ADC) cytotoxicity assays and the development of chimeric antigen receptor (CAR) T cells targeting CDH3, tested in prostate cancer cell lines and xenograft models.

**Results:** *Pten/Apc* double deletion in prostate GEMMs led to highly aggressive tumors with markedly elevated CDH3 expression and enrichment of non-canonical WNT signaling components. Transcriptomic analyses of patient-derived prostate tumors confirmed that CDH3 is significantly upregulated in basal-like prostate cancer subtypes relative to luminal subtypes. Single-cell RNA sequencing revealed CDH3 expression predominantly in basal epithelial cells. Mechanistically, we found that active YAP1 signaling and a WNT5A-ROR2 non-canonical WNT axis drive CDH3 expression. Targeting CDH3 with a CDH3-specific ADC induced potent, antigen-dependent killing of CDH3⁺ prostate cancer cells in vitro and significantly suppressed tumor growth in *in vivo* metastatic prostate cancer models. Likewise, CDH3-targeted CAR T cells specifically recognized and lysed CDH3-expressing prostate tumor cells while sparing CDH3-negative cells, leading to tumor regression and improved survival in mouse models, especially when combined with PD-1 checkpoint blockade.

**Conclusions:** CDH3 is a key marker and functional driver of basal-like prostate cancer. Therapeutic strategies leveraging CDH3, including ADCs and CAR T cells, demonstrate strong preclinical efficacy, supporting the development of CDH3-targeted treatments to overcome resistance in aggressive prostate cancer.

## INTRODUCTION

Prostate cancers can be classified into luminal and basal molecular subtypes that differ markedly in clinical behavior. Luminal-subtype tumors typically respond to androgen deprivation therapies, whereas basal-subtype tumors often exhibit primary resistance to hormonal treatments ^1,2^. Basal-like prostate cancers – especially those arising in castration-resistant disease that lack androgen receptor (AR) activity and neuroendocrine features (so-called “double-negative” prostate cancers) – are associated with aggressive progression and poor outcomes. These observations underscore an urgent need for novel therapeutic targets tailored to treatment-resistant, basal-like prostate cancer.

Recent multi-omics studies have begun to elucidate key pathways driving aggressive prostate tumors, highlighting dysregulated stem-like programs ^3^, aberrant WNT signaling (for example, through *APC* loss) ^4,5^, and alterations in cell-adhesion molecules such as cadherins ^6,7^. Among the cadherin family, CDH3, encoded by *CDH3*, has emerged as a candidate oncogenic driver in multiple solid tumors including breast and prostate cancers ^8^. In normal tissues, CDH3 expression is largely restricted to specific epithelial compartments. In contrast, it is frequently overexpressed in tumor cells with basal-like or stem-like phenotypes – populations linked to enhanced metastatic capacity, therapeutic resistance, and poor prognosis ^9–11^. These findings suggest that CDH3 might contribute to the aggressive nature of basal-like prostate cancer.

Despite mounting evidence implicating CDH3 in cancer progression, there are currently few therapeutic approaches targeting this protein, particularly in prostate cancer. Two promising immunotherapeutic modalities are antibody–drug conjugates (ADCs) and chimeric antigen receptor (CAR) T cells. ADCs use tumor-selective antibodies to deliver cytotoxic payloads into cancer cells and have shown efficacy in various malignancies ^12,13^. CAR T cells harness the patient’s T cells, engineered with synthetic receptors, to recognize and destroy cancer cells with high specificity ^14–16^. Both strategies require an antigen that is abundantly and selectively expressed on tumor cells but minimally present in normal tissues. CDH3 is a compelling candidate in this regard, given its elevated expression in basal-like tumors and limited distribution in most normal adult tissues.

In the present study, we integrate evidence from *in vivo* mouse models, patient tumor datasets, and *in vitro* functional assays to establish CDH3 as a pivotal marker and therapeutic target in basal-like (double-negative) prostate cancer. We demonstrate that loss of *Pten* and *Apc* in the prostate drives a CDH3-high, WNT-active tumor state reminiscent of human basal-like prostate cancer. We further show that CDH3 expression is sustained by YAP1 and non-canonical WNT signaling, and that interrupting this axis can downregulate CDH3. Finally, we provide proof-of-concept that targeting CDH3 with a CDH3-specific ADC or CAR T cells yields potent anti-tumor effects in preclinical prostate cancer models. Collectively, our findings pave the way for CDH3-directed therapies to improve outcomes in aggressive prostate cancer.

## RESULTS

### CDH3 Upregulation Accompanies WNT Pathway Dysregulation in Pten/Apc Double-Knockout Mouse Prostate Tumors

To investigate the contribution of *APC* inactivation to prostate tumorigenesis, we first surveyed prostate cancer genomics data for alterations in key tumor suppressor genes. Across multiple patient cohorts queried via cBioPortal ^17^, *APC* was among the most frequently mutated or deleted tumor suppressors (alongside *PTEN, TP53,* and *SMAD4*), suggesting that loss of APC might cooperate with other driver events during prostate cancer progression. This prompted us to generate a series of prostate-specific genetically engineered mouse models using *PB-Cre4* to delete *Pten^L/L^* alone or in combination with other tumor suppressors (such as *Apc ^L/L^, Trp53 ^L/L^,* or *Smad4 ^L/L^*). Notably, prostate cancer models with single *Pten* loss, or double losses like *Pten/p53* or *Pten/Smad4*, have been well established in prior studies ^18–20^. In our cohort, mice with double knockout of *Pten* and *Apc* (PB-Cre; *PtenL/LApcL/L*) developed significantly more aggressive tumors than mice with *Pten* loss alone or *Pten/p53* double loss. *Pten/Apc*–deficient animals showed shortened overall survival and accelerated tumor progression (median survival 24.4 weeks, vs. much longer latency in *Pten* single-knockouts; **Figure 1A**). Histologically, by 4 months of age the prostates of *Pten/Apc* double-knockout mice exhibited invasive adenocarcinoma, whereas age-matched *Pten*-only mutants displayed mostly high-grade prostatic intraepithelial neoplasia (**Figure 1B**). The *Pten/Apc* lesions also manifested early metastasis: using the mTmG fluorescent lineage reporter, we readily detected GFP⁺ disseminated tumor cells or micro-metastases in lung in *Pten/Apc* mice (**Figure S1B**). These findings indicate the tumor-promoting effect of *Apc* loss in the context of *Pten* deletion, driving metastatic progression.

**Fig. 1.**
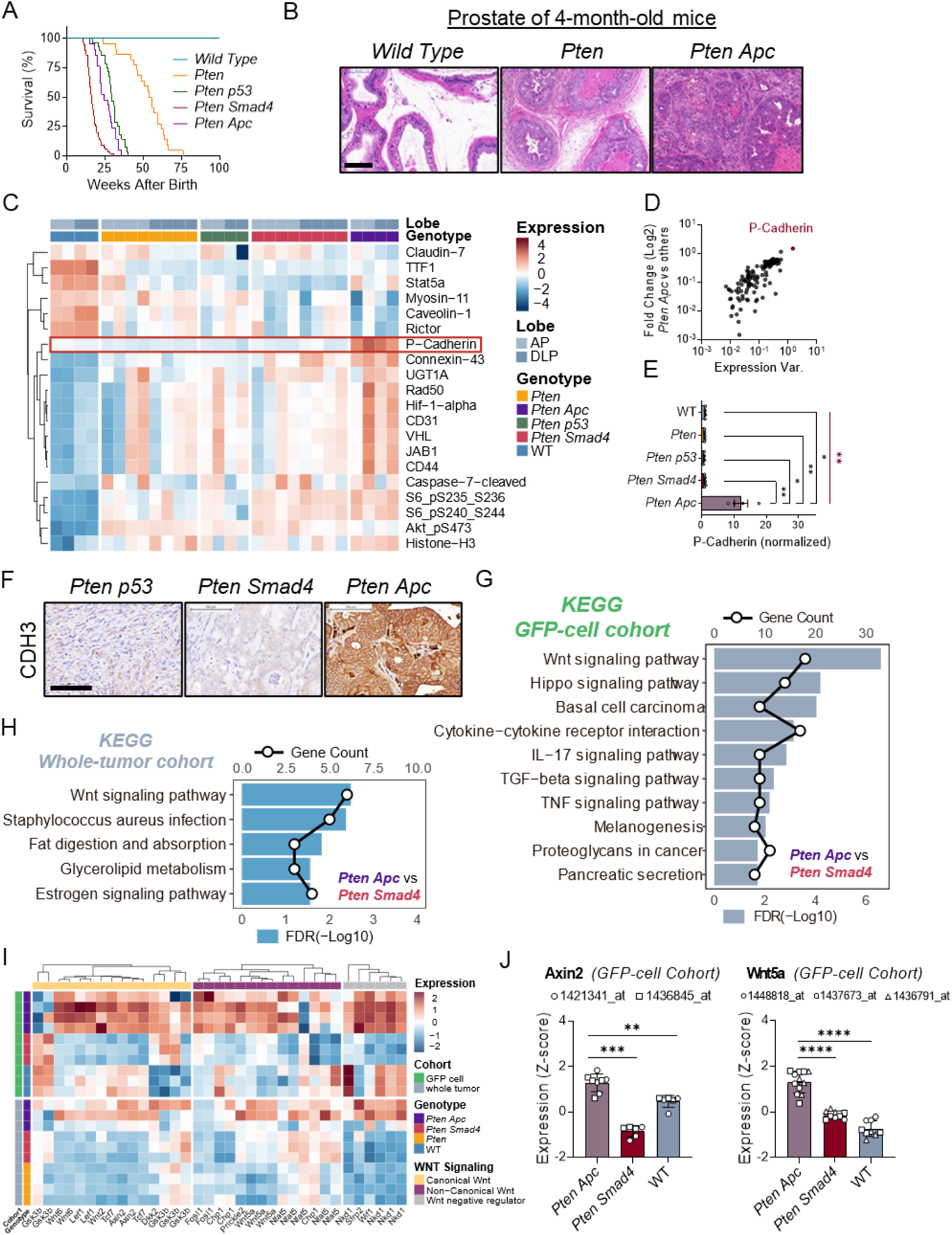
Cdh3 upregulation and WNT pathway dysregulation in the *Pten*/*Apc* prostate specific knockout mouse. (**A**) Survival curves of wild-type (WT) mice compound transgenic mice carrying prostate-specific Cre driver (*PB^Cre^*), floxed fluorescent Cre-reporter (*ROSA^mT/mG^*) and the floxed alleles of key tumor suppressor genes (*Pten* alone or in combination with *p53*, *Smad4* or *Apc*). (**B**) H&E staining of prostate tissue from wild type, *Pten* and *Pten Apc* mice at 4 months old. Scale bar: 100 µm. (**C**) Heatmap of RPPA analysis depicting the top 20 most variably expressed proteins from lysates of wild type prostate and four different genotypes (*Pten*, *Pten Apc*, *Pten p53*, *Pten Smad4*), sampled from the anterior prostate (AP) or dorsolateral prostate (DLP) lobes. The intensity scale ranges from downregulated (blue) to upregulated (red). (**D**) Scatter plot showing the log2 fold change of protein expression (*Pten Apc* vs others) versus expression variability across 281 proteins in all samples. CDH3, the protein with the highest differential expression, is highlighted in red. (**E**) Quantification of CDH3 expression levels (normalized values) across the same five genotypes. (**F**) Immunohistochemical staining showing CDH3 expression in *Pten Apc*, *Pten p53* and *Pten Smad4* tumors. Scale bar: 100 µm. (**G**) KEGG pathway analysis of GFP+ tumor cell transcriptomes (*Pten Apc* vs. *Pten Smad4*) using genes significantly upregulated (log2FC > 2.5, adj. p-value < 0.05). (**H**) KEGG pathway analysis of bulk tumor transcriptomes (*Pten Apc* vs. *Pten Smad4*) using genes significantly upregulated (log2FC > 2, adj. p-value < 0.05). (**I**) Heatmap of the top differentially expressed Wnt pathway–related genes in both the whole-tumor and GFP+ cohort transcriptomes, separated by canonical (yellow labels), non-canonical (purple labels) Wnt signaling and Wnt negative regulators (gray labels). Genes were selected based on differential expression (*Pten Apc* vs. *Pten Smad4*, |log₂FC| > 2, adj. p-value < 0.05) in the GFP⁺ cohort. (**J**) Expression levels of *Axin2* (canonical Wnt marker) and *Wnt5a* (non-canonical Wnt marker) in the GFP+ cell cohort from wild type prostate, *Pten Apc*, and *Pten Smad4* tumors, multiple probe IDs are shown for each gene. Error bars represent SEM. Kruskal-Wallis test (non-parametric ANOVA, red asterisks) for (**D**) and Mann-Whiney test (black asterisks) for (**E**) and (**J**). *P < 0.05, **P < 0.01, ***P < 0.001, ****P < 0.0001.

We next characterized molecular changes associated with the aggressive phenotype of *Pten/Apc*–null prostate tumors. Reverse-phase protein array (RPPA) analysis was performed to compare protein expression across prostate tumors from *Pten*, *Pten/p53*, *Pten/Smad4*, and *Pten/Apc* mice. Among the top differentially upregulated proteins in *Pten/Apc* tumors was CDH3 (**Figure 1C-D**). Quantitative analyses confirmed that CDH3 protein levels were dramatically higher in the *Pten/Apc* group than in tumors from the other genotypes (**Figure 1E**). Immunohistochemistry further validated robust CDH3 expression specifically in *Pten/Apc* prostate tumors, whereas *Pten/p53* and *Pten/Smad4* lesions showed little or no staining (**Figure 1F**). These data identify CDH3 as a uniquely upregulated protein in the context of combined *Pten* and *Apc* loss.

To expand the gene expression detection, we performed RNA transcriptomics on bulk tumor tissue and on sorted epithelial tumor cells (GFP⁺, from mTmG mice) from *Pten/Apc* and other genotypes (**Figure S1D)**, and conducted pathway enrichment analysis. As expected, *Pten/Apc* tumors exhibited a strong WNT signaling signature relative to *Pten* or *Pten/Smad4* tumors (enriched in KEGG, WikiPathways, and Hallmark gene sets; **Figure 1G-H** and **Figure S1E–S1F**). Closer examination revealed that both canonical WNT signaling components with *Axin2* as a representative and non-canonical WNT signaling components with *Wnt5a* as a representative were elevated in *Pten/Apc* whole tumors and GFP⁺ tumor cells (**Figure 1I-J, Figure S1G**). Given the prominent role of Wnt5a in the non-canonical WNT signaling, we further validated higher Wnt5a protein level in *Pten/Apc* prostate tumors compared with *Pten/Smad4* tumor (**Figure S1H**). These indicate that *Apc* loss at the backdrop of *Pten* loss in mouse prostate activate both canonical and non-canonical branches of WNT pathways ^21,22^.

In summary, the *Pten/Apc* double-knockout model produces highly invasive prostate tumors characterized by markedly elevated CDH3 expression and widespread activation of WNT signaling. This raises the question of whether CDH3 upregulation is merely a bystander effect of aggressive tumor biology or a feature of broader relevance in human prostate cancer, particularly in basal-like subtypes where WNT activity and castration resistance often coincide. We therefore next evaluated CDH3 expression in clinical prostate cancer cohorts with respect to molecular subtypes.

### CDH3 Is Associated with Basal-Like Phenotypes in Advanced Prostate Cancer

To determine the clinical relevance of the CDH3 upregulation observed in our mouse model, we analyzed human prostate cancer datasets. We first re-examined a published transcriptomic study of advanced prostate cancers by Tang *et al.* ^23^. In that study, metastatic castration-resistant prostate cancer (CRPC) samples (including patient-derived xenografts, organoids, and cell lines) were classified into subtypes based on chromatin accessibility profiles: namely, AR-dependent, neuroendocrine (NEPC), WNT-driven, and stem cell-like (SCL) subtypes. Notably, the WNT and SCL groups lack both AR activity and neuroendocrine markers – thus representing “double-negative” prostate cancer (DNPC) subtypes analogous to basal-like tumors. Upon re-analysis of the RNA-seq data from Tang *et al.* (**Figure 2A**), we found that both the WNT and SCL subtype tumors exhibit significantly higher *CDH3* expression compared to AR-driven or NEPC tumors (**Figure 2B-C**). These DNPC subtypes also showed elevated basal epithelial gene signatures and WNT pathway activity, with the WNT subtype biased toward canonical WNT activation and the SCL subtype showing higher non-canonical WNT and stemness signatures (**Figure 2C**, **Figure S2A**). This re-analysis confirms *CDH3* expression in basal-like, AR-independent prostate cancer in human.

**Fig. 2.**
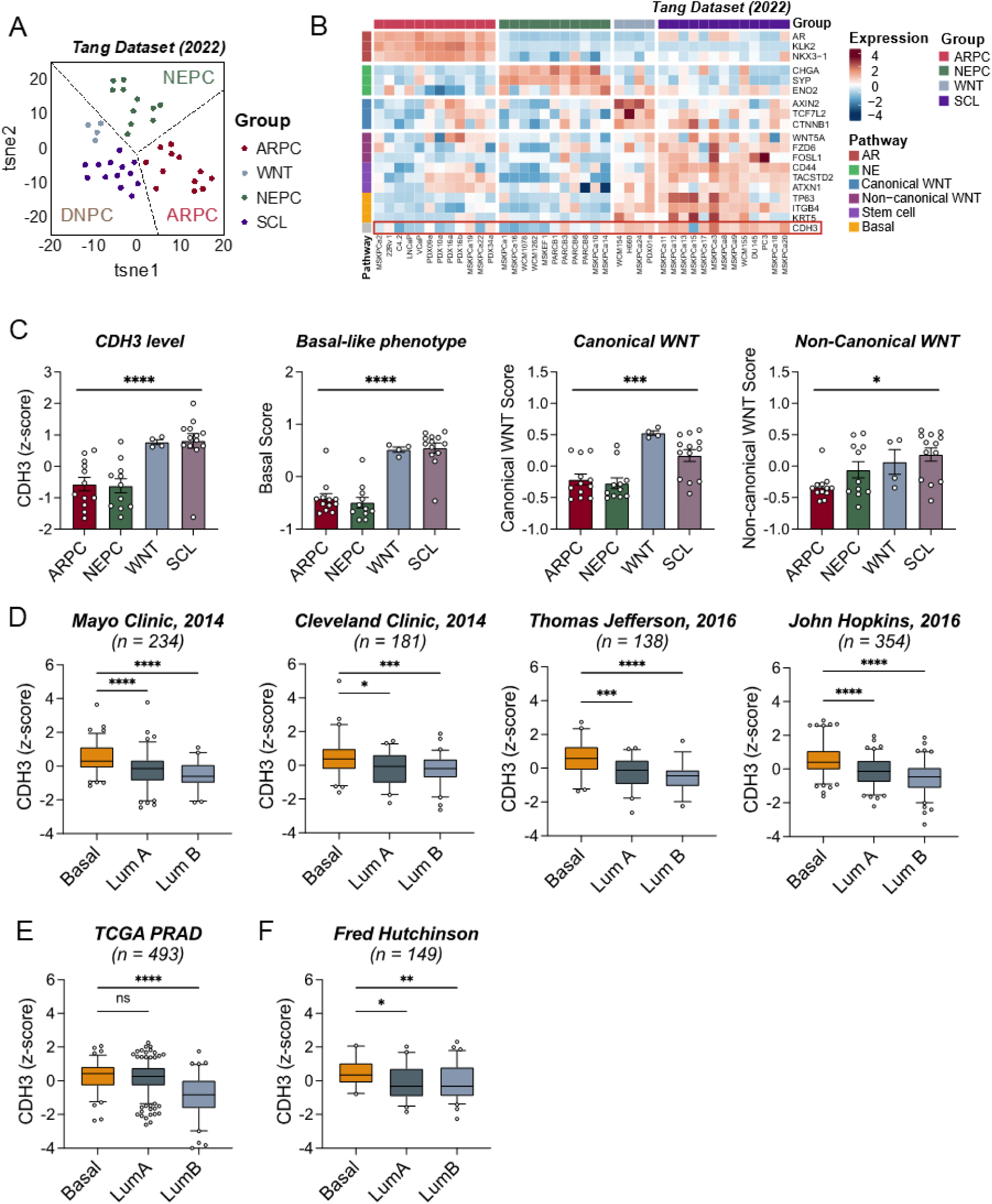
CDH3 is highly correlated basal-like phenotype in CRPC, and elevated in basal-like prostate cancer and cancer. (**A**) t-SNE projection of transcriptomes of CRPC patient-derived xenograft (PDX), cell line, and organoid samples from Tang et al. (2022), illustrating four major subtypes: AR-dependent (ARPC), stem cell-like (SCL), neuroendocrine (NEPC), and WNT-driven (WNT). (**B**) Heatmap of selected pathway-associated genes across the same sample set. Red denotes upregulation, blue denotes downregulation. (**C**) CDH3 expression level, canonical WNT score, non-canonical WNT score, and basal score, compared among the different groups. (**D**) CDH3 expression in different subtype groups across multiple prostate cancer cohorts (Mayo, Cleveland, Thomas Jefferson, Johns Hopkins). Samples were classified into basal-like, luminal A, or luminal B subtypes using a PAM50-based approach. (**E**) CDH3 expression in TCGA primary prostate cancer (PRAD), classified by PAM50 subtypes, (**F**) Metastatic sample from Fred Hutch Cancer Center (FHCRC) prostate cancer datasets, classified by PAM50 subtypes. Error bars represent SEM. Kruskal-Wallis test (non-parametric ANOVA) for (**C**) and Mann-Whiney test for (**D**), (**E**) and (**F**). Box plots represent the 5th to 95th percentiles, with the horizontal line indicating the median for (**D**), (**E**) and (**F**). *P < 0.05, **P < 0.01, ***P < 0.001, ****P < 0.0001, ns, not significant.

We extended our investigation to clinical prostate tumor cohorts stratified by the PAM50 intrinsic subtyping scheme. PAM50, originally developed for breast cancer, classifies tumors into molecular subtypes (Luminal A, Luminal B, HER2-enriched, Basal-like) based on gene expression profiles ^24^. Although prostate cancer is not typically classified by PAM50 in the clinic, recent work has applied this approach to prostate tumors ^25^. We applied PAM50 classification to multiple prostate cancer gene expression cohorts (including datasets from the Mayo Clinic ^26^, Cleveland Clinic ^27^, Thomas Jefferson Hospital ^28^, and Johns Hopkins University ^29^). Across all these cohorts, we consistently observed that tumors classified as basal-like expressed significantly higher *CDH3* levels than those classified as luminal A or luminal B (**Figure 2D**). A similar pattern was evident in the TCGA Prostate Adenocarcinoma (PRAD) dataset, where basal-like cases had greater *CDH3* expression compared to luminal cases (**Figure 2E**). Furthermore, in a cohort of metastatic prostate tumors from the Fred Hutchinson Cancer Research Center, basal-type metastases had the highest *CDH3* expression (**Figure 2F**).

Together, these analyses of human data demonstrate that *CDH3* overexpression is a hallmark of basal-like prostate cancer. Tumors with basal or double-negative features – whether defined by lack of AR/NE markers or by PAM50 subtyping – tend to have elevated CDH3 levels.

### CDH3 Is Preferentially Expressed in Basal Epithelial Cells Alongside Non-Canonical WNT Signaling

Given the strong association of CDH3 with basal-like prostate tumors, we sought to pinpoint the cellular context of CDH3 expression within tumors. To explore this, we re-analyzed published single-cell RNA sequencing (scRNA-seq) data of human prostate cancer ^30 31^. Each scRNA-seq dataset was annotated for major cell lineages – epithelial tumor cells, immune cells, and stromal cells – using canonical markers (e.g., *NKX3-1* and *KRT18* for luminal prostate cells, *KRT5* and *KRT14* for basal cells, *CD45/PTPRC* for immune cells). We then examined *CDH3* expression within the epithelial compartment, further distinguishing basal-like from luminal-like epithelial tumor cells based on their gene signatures.

Song *et al.* dataset contained most major cell types in solid tumors, including basal and luminal epithelial cell clusters (**Figure 3A**, **Figure S3A**). *CDH3* was predominantly expressed in basal-type epithelial cells, with minimal expression in luminal tumor cells (**Figure 3B**, only showing epithelial cells). Moreover, a few key genes involved in non-canonical WNT signaling, including *WNT5A*, *ROR2*, and downstream transcription factor *FOSL1* (an AP-1 family member), were co-enriched in the basal epithelial population (**Figure 3C**). In contrast, classical canonical WNT pathway components (e.g., *CTNNB1* encoding β-catenin, *LRP5/6* encoding co-receptors) did not show a strong preference for basal over luminal cells (**Figure 3D**). Equivalent results were obtained from Wong et al. dataset (**Figure 3E-H, Figure S3B**), together suggesting that CDH3⁺ basal cells are associated with an active non-canonical WNT signaling in prostate tumors.

**Fig. 3.**
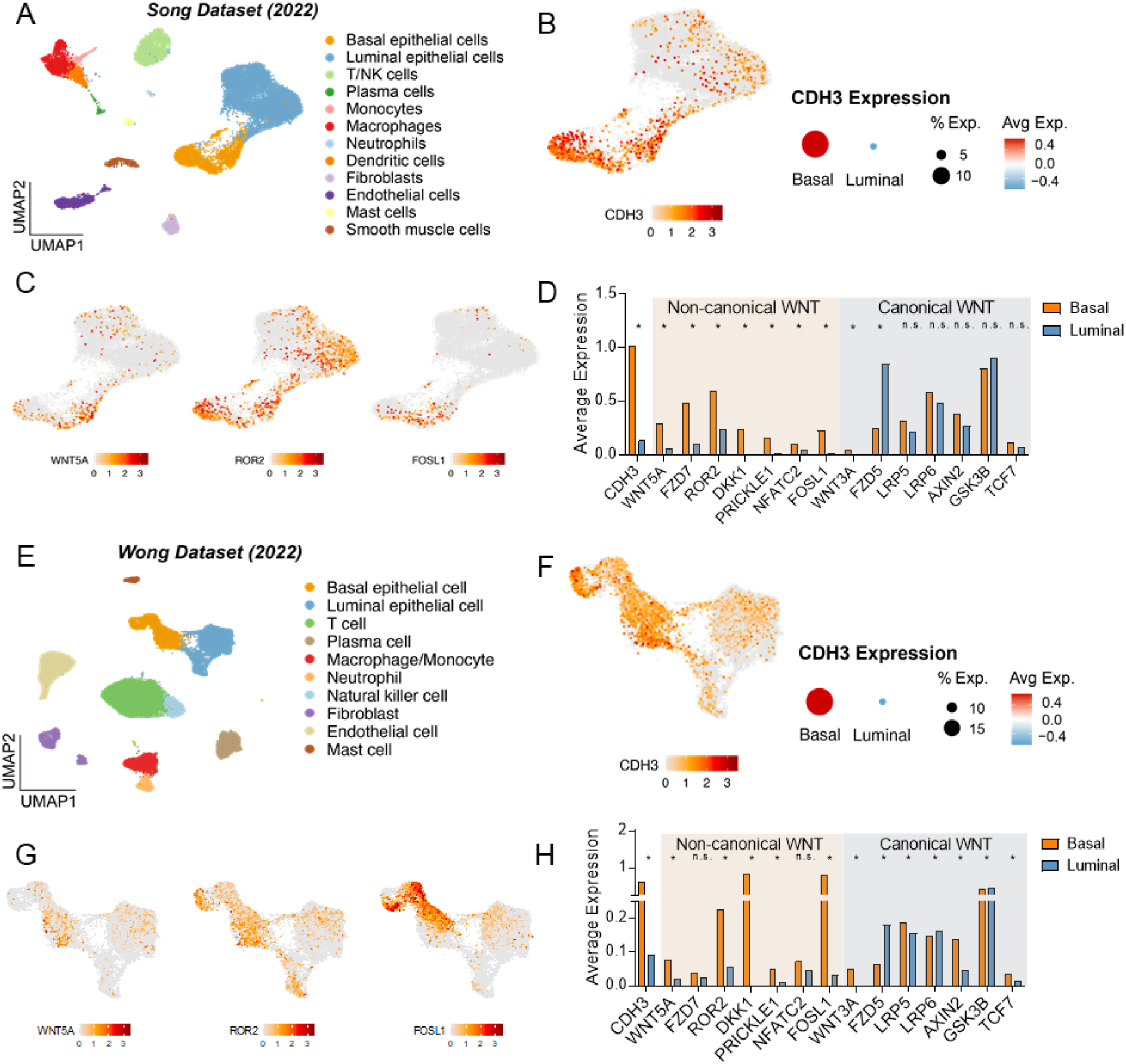
CDH3 has Preferential Association with Non-canonical WNT signaling in basal cells. (**A**) Single-cell UMAP from the Song et al. (2022) dataset showing prostate cancer cells grouped into basal epithelial, luminal epithelial, immune, and stromal populations. (**B**) CDH3 expression in the epithelial sub-compartment in feature plot (heatmap scale) and dot plot (dot size indicates percentage of expressing cells; color represents average expression). (**C**) Feature plots for selected non-canonical WNT genes within the epithelial sub-compartment. (**D**) Comparison of average expression of canonical vs. non-canonical WNT pathway components in basal vs. luminal cells. (**E**) Single-cell UMAP from the Wong et al. (2022) dataset, illustrating a similar array of cell populations. (**F**) CDH3 expression in the epithelial sub-compartment in feature plot (heatmap scale) and dot plot (dot size indicates percentage of expressing cells; color represents average expression). (**G**) Feature plots for selected non-canonical WNT genes within the epithelial sub-compartment. (H) Comparison of average expression of canonical vs. non-canonical WNT pathway components in basal vs. luminal cells. Wilcoxon rank-sum test for (**D**) and (**H**). *Adjusted P < 0.01, ns, not significant.

We obtained similar insights from an additional scRNA-seq dataset that included castration-resistant and neuroendocrine prostate cancer cells (Jee et al. dataset, GSE221603). There too, *CDH3* expression and non-canonical WNT markers were both preferentially elevated in basal-like epithelial cells (**Figure S3C-E**), implying a coupling between CDH3 and WNT5A/ROR2 signaling in those cells. Collectively, CDH3 expression is confined largely to the basal epithelial compartment of prostate tumors, where it correlates with non-canonical WNT activity, prompting us to explore whether CDH3 expression is regulated by non-canonical WNT pathway.

### CDH3 Expression is Regulated by YAP1 through a WNT5A–ROR2 Non-Canonical WNT Axis

The convergence of YAP1 signaling, non-canonical WNT signaling, and CDH3 expression in basal-like prostate cancer led us to investigate whether these pathways interact functionally. Prior research indicated that YAP (Yes-associated protein), a Hippo pathway effector often active in aggressive prostate cancers, can cooperate with AP-1 transcription factors like FOSL1 to promote a stem-like, castration-resistant tumor state (the SCL subtype) ^23^. Additionally, recent studies reported that in prostate cancer, YAP activity can induce *WNT5A* and *ROR2* expression, and conversely, WNT5A–ROR2 signaling can modulate YAP as a feedback mechanism ^32^. We therefore hypothesized that YAP might drive *CDH3* expression via non-canonical WNT signaling components WNT5A and ROR2.

To test this, we first examined a panel of human prostate cancer cell lines with varying CDH3 levels: DU145 and 22Rv1 with high CDH3, LNCaP and LHMK with intermediate/low CDH3, and PC3 with low CDH3. In CDH3-high lines (DU145, 22Rv1), YAP1 was predominantly localized in the nucleus indicative of active function, whereas in CDH3-intermediate/low lines (LNCaP, PC3, LHMK), YAP1 was mostly cytoplasmic and inactive (**Figure 4A**). Consistent with the subcellular localization, CDH3-high cell lines DU145 and 22Rv1 had the lower ratios of phosphor-YAP/total-YAP than the CDH3-low cell lines when assessed with western blot (**Figure S4A–S4B**). These observations uphold a positive role of YAP activity in regulating CDH3 expression in prostate cancer cells. We then directly inhibited YAP to assess effects on *CDH3*. Treatment of DU145 (CDH3-high) cells with verteporfin, a YAP-TEAD interaction inhibitor, re-localized YAP from nucleus to cytoplasm and caused a dose-dependent decrease in CDH3 expression (**Figure 4B–C**, **Figure S4C**), showing that active YAP signaling is required to maintain high CDH3 expression in these cells.

**Fig. 4.**
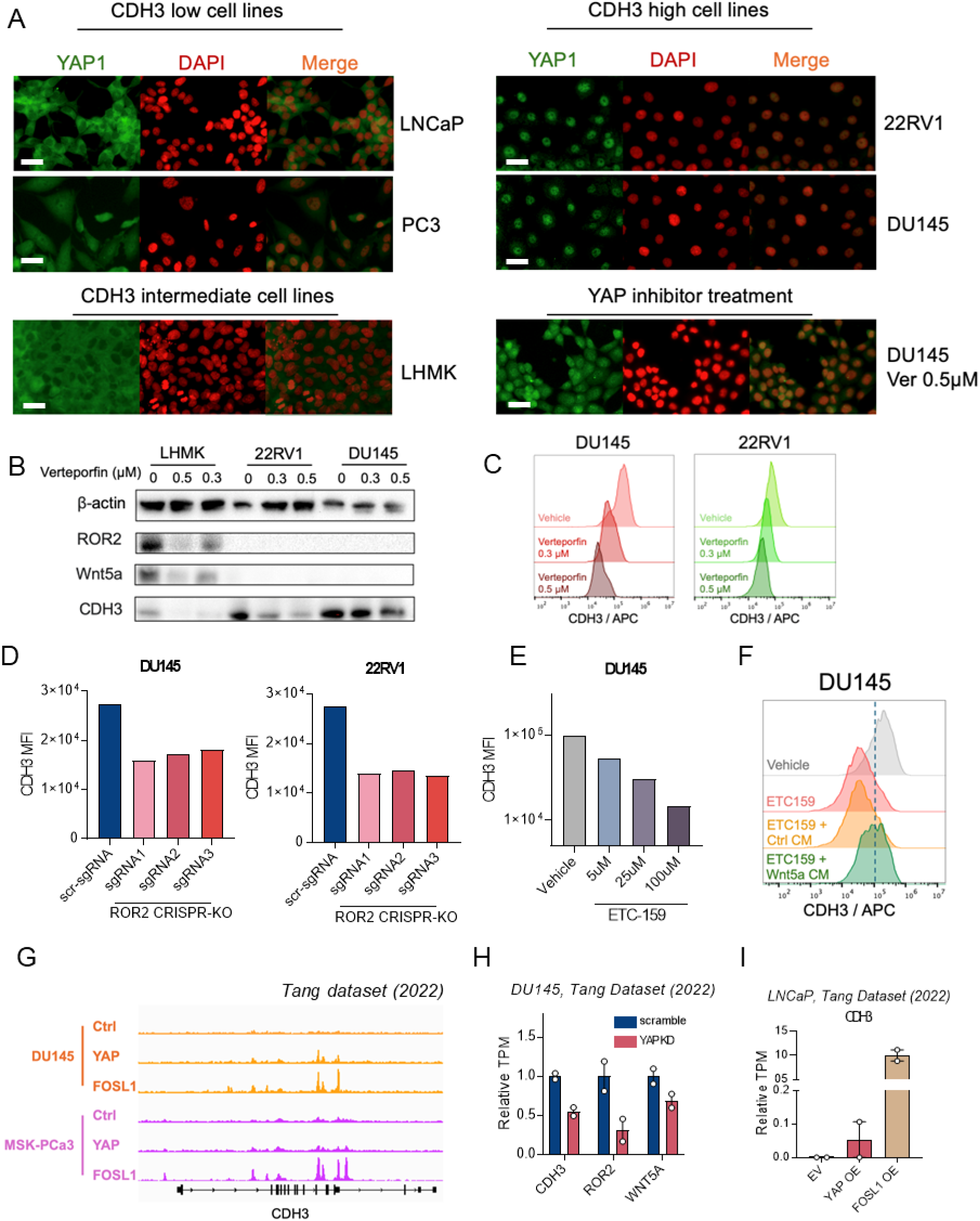
CDH3 regulation through YAP/ WNT5A / ROR2. (**A**) Immunofluorescence of prostate cancer cell lines (LNCaP, PC3, LHMK, and DU145 cells), YAP1 (green) and nuclei (DAPI, red) are shown to illustrate subcellular localization. An extra group of DU145 cells were treated with the YAP inhibitor verteporfin (Ver, 0.5 µM) for 72 hours. Scale bar, 20µm. (**B**) Western blot of CDH3 protein levels in LHMK, 22RV1, and DU145 cells comparing vehicle controls to verteporfin-treated conditions (0.3, 0.5 µM) for 72 hours. β-actin serves as a loading control. (**C**) Flow cytometry histograms of CDH3 expression in DU145 and 22RV1 cells, comparing vehicle controls to verteporfin-treated conditions (0.3, 0.5 µM) for 72 hours. (**D**) Flow cytometry analysis of CDH3 median fluorescence intensity (MFI) in DU145 and 22RV1 cells following CRISPR-mediated ROR2 knockout (sgRNA1–3), compared to a non-targeting control (scr-sgRNA). (**E**) Flow cytometry analysis of CDH3 MFI in DU145 cells, comparing vehicle controls to ETC159 (5, 25, 100 μM, an inhibitor of WNT ligand secretion) conditions for 72 hours. (**F**) Flow cytometry histograms of CDH3 expression in DU145 cells treated with ETC159 in the presence or absence of Wnt5a-conditioned medium (CM, 20% v/v) for 72h. (**G**) Visualization of CHIP-seq data from Tang et al. (2022) illustrating control (Ctrl), YAP, and FOSL1 binding profiles at the CDH3 locus in DU145 (orange) cell line and MSK-PCa3 (purple) organoid. (**H**) Expression levels of CDH3, ROR2, and WNT5A in DU145 cells upon YAP knockdown (KD) versus scramble control. (**I**) Expression levels of CDH3 upon overexpression (OE) of YAP or FOSL1 in LNCaP cells. Error bars represent SEM.

Next, we interrogated the role of the non-canonical WNT pathway. Through analysis of TCGA prostate cancer data, we found that *ROR2* (a receptor for WNT5A) is among the genes most strongly co-expressed with *CDH3* in primary tumors (**Figure S4D-S4E**). We therefore tested whether disrupting WNT5A–ROR2 signaling would impact CDH3. CRISPR/Cas9 knockout of *ROR2* in DU145, 22Rv1 and LHMK cells decreased CDH3 protein levels (**Figure 4D**, and **Figure S4F–S4G**), supporting a requirement for ROR2 in sustaining *CDH3* expression. Moreover, treating CDH3-expressing lines with ETC-159, a small molecule that broadly inhibits WNT ligand secretion, led to a dose-dependent decrease in surface CDH3 (**Figure 4E**, **Figure S4H**). Conversely, adding exogenous WNT5A partially rescued the loss of CDH3 caused by ETC-159 (**Figure 4F**), indicating that WNT5A is both necessary and sufficient to drive CDH3 expression.

To connect these findings with YAP, we leveraged published data from Tang *et al.*’s CRPC study ^23^. ChIP-seq profiles in prostate cancer models indicated that FOSL1 (an AP-1 partner of YAP/TEAD) binds strongly to regulatory regions at the *CDH3* locus, whereas YAP1 itself shows weaker binding (suggesting YAP might act via FOSL1 or other partners) (**Figure 4G**). Tang et al. silenced YAP1 in DU145 cells using RNAi, which led to reduced transcript levels of *CDH3*, *WNT5A*, and *ROR2* (**Figure 4H**). Conversely, overexpressing FOSL1 in CDH3-low LNCaP cells dramatically upregulated *CDH3*, whereas overexpressing YAP1 had a more limited effect, consistent with YAP presumably needing a co-factor like AP-1 to drive *CDH3* transcription (**Figure 4I**). These results place YAP1 upstream of the WNT5A–ROR2–CDH3 circuit: active YAP (together with AP-1) induces WNT5A and ROR2, which in turn help sustain CDH3 expression.

### CDH3-Targeted ADC Therapy Shows Efficacy in Prostate Cancer Models

Given the strong expression of CDH3 in basal-like prostate cancer, we evaluated the immunotherapeutic potential of targeting CDH3. First, we aimed to develop antibody–drug conjugate (ADC) targeting CDH3. We used a humanized IgG1 antibody directed against the extracellular domains of CDH3 (Perseus Proteomics), and chose to focus on DU145 to test the therapeutic agents because of its highest surface CDH3 among the various prostate cancer cell lines (**Figure 5A**). Using CRISPR/cas9, we generated CDH3 knockout (KO) in DU145 (**Figure S5A**) and subsequently labeled the DU145 sublines with a triple reporter (TR) of the HSV1-tk-GFP-luciferase fusion protein^33^. Wild-type DU145 (CDH3^WT^) and knockout (CDH3^KO^) cells were incubated with a pH-sensitive fluorescent anti-CDH3 antibody conjugate (pHrodo), which fluoresces upon internalization into acidic endosomes. CDH3^WT^ cells showed robust dose-dependent internalization of the antibody, visualized as punctate intracellular fluorescence, whereas CDH3^KO^ cells displayed minimal uptake (**Figure 5B**). CDH3-targeted ADC was generated by conjugating anti-CDH3 IgG1 to a thioester linker-maytansinoid payload (DM1) at 3∼4 DM1 warheads per IgG (**Figure 5C**). Flow cytometry quantification with a pHrodo-labeled CDH3-ADC similarly showed efficient internalization in DU145-CDH3^WT^ cells but not in DU145-CDH3^KO^ cells (**Figure S5B-C**). These data confirm that CDH3 on tumor cells mediates antibody binding and internalization – an essential property for ADC function.

**Fig. 5.**
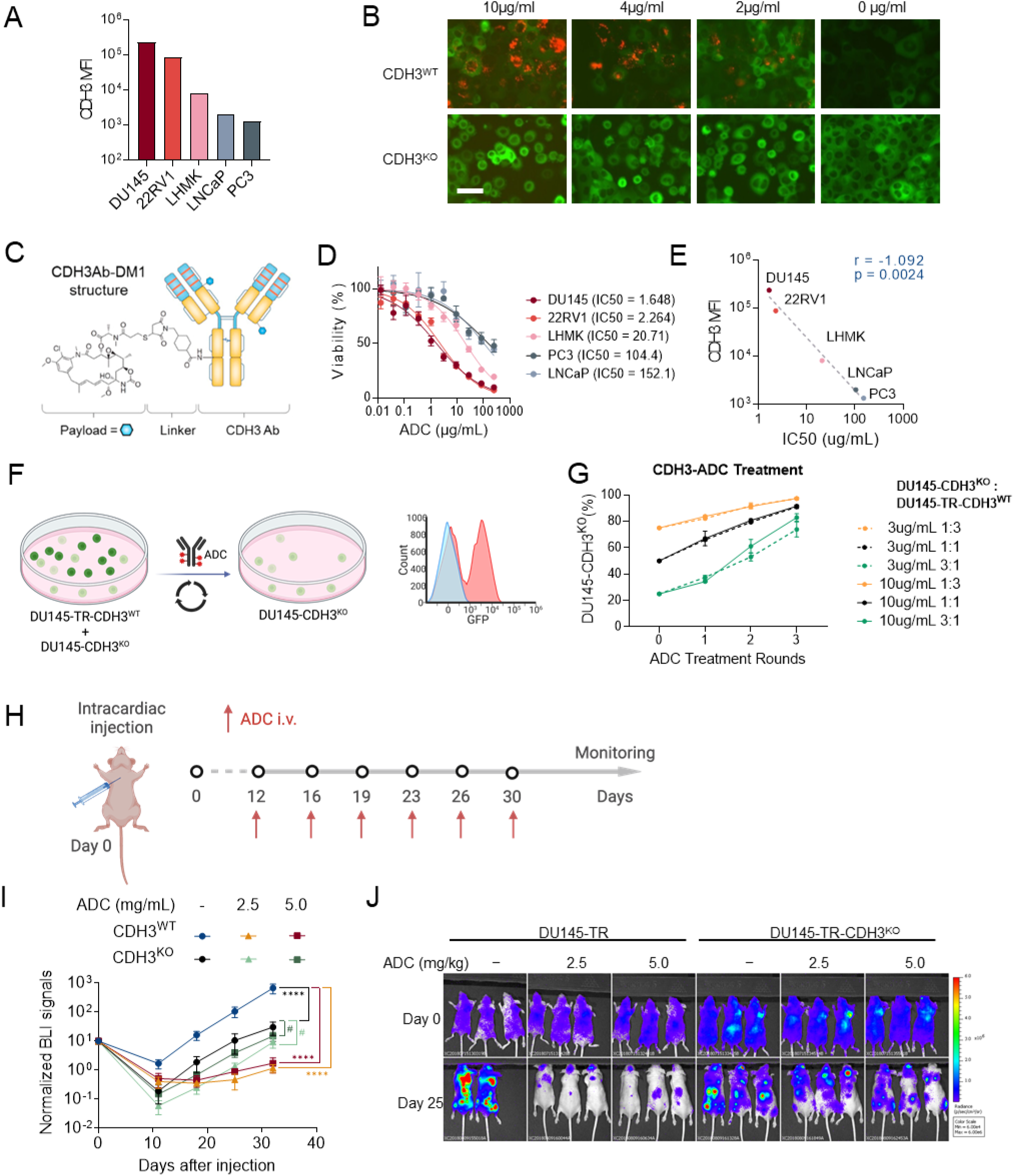
CDH3 targeted ADC treatment. (**A**) Flow cytometry analysis of CDH3 MFI in different prostate cancer cell lines (DU145, 22RV1, LHMK, LNCaP, PC3). (**B**) Immunofluorescence of DU145-TR-CDH3^WT^ (top) and DU145-TR-CDH3^KO^ (bottom) cells treated with different concentrations of CDH3 antibody (10, 4, 2, and 0 µg/mL) for 48 hours. Green fluorescence (TR-labeled cells) represents the cells, and red fluorescence (pHrodo-labeled antibody) indicates antibody internalization. Scale bar, 50µm. (**C**) Schematic representation of the CDH3Ab–DM1 antibody–drug conjugate, illustrating the anti-CDH3 antibody, linker, and cytotoxic DM1 payload. (**D**) Dose-dependent viability of five prostate cancer cell lines (DU145, 22RV1, LHMK, PC3, LNCaP) treated with the CDH3Ab–DM1 conjugate with their respective IC50 values. (**E**) Correlation between CDH3 surface expression (MFI) and CDH3Ab-DM1 IC50 across multiple cell lines, with correlation coefficient (r) and significance (p) shown. (**F**) Schematic illustrating the co-culture setup of DU145-TR (CDH3 wild type, GFP-labeled) and DU145-CDH3 knockout (non-labeled) cells, followed by cyclical ADC treatments at varying concentrations and different initial mixing ratios. Red and blue histograms indicate the expected result before and after treatment, respectively. (**G**) The proportion of DU145-CDH3^KO^ cells over three rounds of ADC treatment, comparing 3 µg/mL and 10 µg/mL doses across different initial co-culture ratios (DU145-TR-CDH3^WT^ : DU145-CDH3^KO^ at 1:3, 1:1 and 3:1). (**H**) Schematic of intracardiac injection and intravenous ADC administration schedule for nude mice bearing DU145-TR or DU145-TR-CDH3^KO^ cells, with treatment days indicated by red arrows. (**I**) Normalized bioluminescent intensity (BLI, scaled to Day 0 as 10) signals measured in legs over 30 days in mice injected with DU145-TR-CDH3^WT^ or DU145-TR-CDH3^KO^ cells. N=10. (**J**) Bioluminescent imaging of DU145-TR-CDH3^WT^ or DU145-TR-CDH3^KO^ tumor burden at Day 0 and Day 25 under different ADC doses (2.5 mg/kg or 5 mg/kg) versus untreated controls. Error bars represent SEM. Mann-Whiney test for (**I**). *P < 0.05, **P < 0.01, ***P < 0.001, ****P < 0.0001, #, not significant.

We then tested this ADC on a panel of prostate cancer cell lines (DU145, 22Rv1, LHMK, PC3, LNCaP). In vitro cytotoxicity assays demonstrated that the ADC caused potent, dose-dependent killing of CDH3-positive cells. Cell lines with higher CDH3 expression were far more sensitive, showing low IC₅₀ values, whereas low-CDH3 lines like LNCaP were relatively resistant (**Figure 5D**). There was a strong inverse correlation between surface CDH3 level and ADC IC₅₀ (**Figure 5E**), indicating that antigen density largely governs ADC effectiveness.

To further confirm that the cytotoxicity is specific to CDH3-expressing cells, we performed a mixed-culture competition assay. We co-cultured GFP^+^ DU145-TR-CDH3^WT^ cells and unlabeled DU145-CDH3^KO^ cells at defined ratios, then treated the mixture with the ADC over multiple cycles (**Figure 5F**). Over the course of three treatments, the fraction of CDH3^WT^ cells in the population declined drastically, indicating selective elimination of CDH3^+^ cells by the ADC (**Figure S5D)**. In contrast, untreated control co-cultures maintained a stable mix of WT and knockout cells (**Figure S5E)**. By the end of treatment, CDH3^KO^ cells had overgrown the culture, having effectively been “spared” by the ADC (**Figure 5G**). This competition experiment underscores the specificity of the ADC for CDH3^+^ cells.

Finally, we evaluated the therapeutic impact of the CDH3-ADC in vivo using prostate cancer xenograft models. We employed an aggressive metastatic model by intracardiac inoculation of DU145-TR cells. Two groups of immunodeficient mice were injected with either DU145-TR-CDH3^WT^ or DU145-TR-CDH3^KO^, establishing systemic tumor colonization (**Figure 5H**). Once metastases were evident, mice were treated with the CDH3-ADC or vehicle control. In mice bearing CDH3^WT^ tumors, ADC administration (2.5 mg/kg or 5 mg/kg IV twice/weekly) led to a significant reduction in metastatic tumor burden as measured by bioluminescence imaging, and extended survival relative to control-treated mice (**Figure 5I-J**). In contrast, mice harboring CDH3^KO^ tumors did not response to the ADC (**Figure 5I-J**). These in vivo results confirm that CDH3-targeted ADC therapy can effectively suppress metastatic prostate tumor growth, provided the tumor expresses sufficient CDH3. Together, the evidence supports the potential of a CDH3-directed ADC to selectively eliminate CDH3–expressing prostate cancer cells in vivo.

### CDH3-Targeted CAR T Cells Mediate Specific Tumor Cell Killing In Vitro

While ADCs can deliver potent cytotoxic drugs to tumors, their efficacy may be limited by dose-dependent systemic toxicities and potential resistance mechanisms^34,35 36^. CAR T cell therapy offers a complementary approach, with the capability for active tumor surveillance and long-term persistence of effector cells ^14–16^. We therefore developed CAR T cells targeting CDH3 to assess whether T-cell mediated attack could likewise exploit CDH3 as a vulnerability in basal-like prostate cancer.

We constructed a second-generation CAR bearing a CDH3-specific single-chain variable fragment (scFv) derived from the same antibody used in our ADC. The CAR included a CD8α hinge and transmembrane domain, 4-1BB (CD137) co-stimulatory domain, and CD3ζ signaling domain (**Figure 6A**), in line with established CAR designs ^37,38^. The functionality and specificity of the anti-CDH3 CAR was tested using Jurkat T cells, which were efficiently transduced with the CAR as indicated by the co-expressed GFP (**Figure 6B**). Jurkat transduced with αCDH3-CAR showed robust activation, as measured by elevated CD69 and CD25, when co-cultured with DU145-CDH3^WT^ but not DU145-CDH3^KO^ cells, and Jurkat transduced with an irrelevant CAR (anti-CD19) remained non-responsive (**Figure 6C**, **Figure S6A**). Having verified the CAR, we proceeded to generate αCDH3-CAR T cells from primary human T lymphocytes. Peripheral blood mononuclear cells were transduced with the αCDH3-CAR or αCD19-CAR and expanded *ex vivo* (**Figure 6D)**. After expansion, flow cytometry confirmed that the CAR T cell products had comparable compositions of CD4⁺ and CD8⁺ T cells (∼1:1 ratio) (**Figure S6B**).

**Fig. 6.**
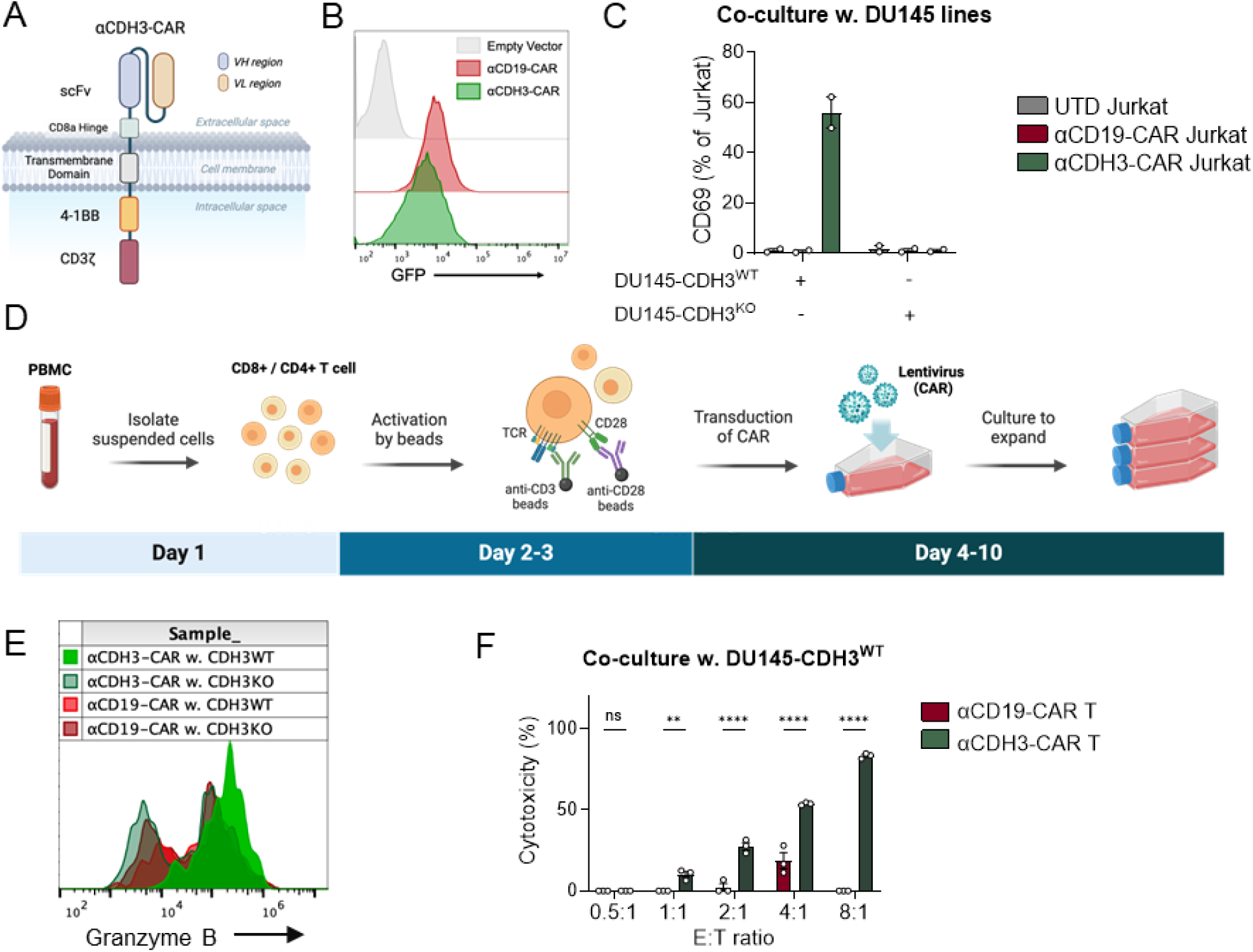
CDH3 targeted CAR T in vitro. (**A**) Schematic diagram of the αCDH3-CAR construct containing scFv, CD8α hinge, transmembrane domain, 4-1BB costimulatory domain, and CD3ζ signaling domain. (**B**) Flow cytometry histograms indicating αCD19-CAR and αCDH3-CAR constructs (both carry a GFP gene) expression in Jurkat cells. (**C**) Flow cytometry–based quantification of Jurkat cell activation (CD69 and CD25 expression) following coculture with DU145-CDH3^WT^ or DU145-CDH3^KO^ cells, comparing the αCDH3-CAR with untransduced (UTD) and αCD19-CAR control. (**D**) Schematic of the T-cell engineering protocol, showing isolation of PBMCs, bead-based activation, lentiviral transduction of CAR, and subsequent expansion. (**E**) Flow cytometry histogram of granzyme B expression in T cells transduced with either αCD19-CAR or αCDH3-CAR, following coculture with DU145-CDH3^WT^ or DU145-CDH3^KO^ cells. (**F**) Cytotoxicity assays of DU145-CDH3^WT^ or DU145-CDH3^KO^ tumor targets at increasing effector-to-target (E:T) ratios, comparing αCD19-CAR T cells and αCDH3-CAR T cells. Error bars represent SEM. Mann-Whiney test for (**F**). *P < 0.05, **P < 0.01, ***P < 0.001, ****P < 0.0001, ns, not significant.

We then evaluated the anti-tumor function of αCDH3-CAR T cells in co-culture assays with prostate tumor cells. Upon co-incubation with DU145-CDH3^WT^ cells, αCDH3-CAR T cells became strongly activated, as evidenced by upregulation of granzyme B and perforin; neither αCD19-CAR T cells nor interaction with DU145-CDH3^KO^ cells altered the granzyme B and perforin signals (**Figure 6E**, **Figure S6C**). Importantly, αCDH3-CAR T cells exhibited potent cytotoxicity against prostate cancer targets: when co-cultured with DU145-CDH3^WT^ cells at various effector-to-target (E:T) ratios, the αCDH3-CAR T cells lysed the majority of tumor cells at moderate to high E:T ratios, whereas αCD19-CAR T cells did not induce significant tumor cell death (**Figure 6F**). This cytotoxic effect was strictly antigen-dependent, as αCDH3-CAR T cells did not kill any DU145-CDH3^KO^ cells (**Figure S6D**). These in vitro findings demonstrate that αCDH3-CAR T cells selectively target and destroy CDH3–expressing prostate tumor cells, fulfilling the specificity and potency criteria needed to move forward to in vivo testing.

### CDH3-Targeted CAR T Cells Curbed Metastasis, Especially in Combination with Checkpoint Blockade

We next evaluated the anti-tumor efficacy of αCDH3-CAR T cells in vivo. First, to examine whether CAR T cells home and infiltrate tumors, we implanted NOD-SCID mice subcutaneously with DU145-CDH3^WT^ or DU145-CDH3^KO^ cells. After tumors were established, mice received a dose of cyclophosphamide followed by infusion of either αCDH3-CAR T cells or αCD19-CAR T cells. One week later, αCDH3-CAR T cells were found accumulated at higher levels in DU145-CDH3^WT^ tumors than DU145-CDH3^KO^ tumors (**Figure S7A-D**). Next, CAR T therapy was tested with the metastasis model, similar to the ADC experiments. CD145-TR cells populated widespread metastasis after intracardiac injection, and the mice were then treated with cyclophosphamide followed by infusion of αCDH3-CAR T, αCD19-CAR T or untransduced (UTD) T cells (**Figure 7A, Figure S7E**). One day after the infusion, both CAR T cells showed similar distributions with highest frequency in the lungs, a phenomenon commonly seen for IV-injected T cells (**Figure S7F**). When T cell subsets were quantified (naïve, central memory, effector memory, effector), both CD8^+^ and CD4^+^ CAR T cells had a large fraction of central memory subsets, a property considered advantageous for immediate and long-term anti-tumor responses (**Figure 7B, Figure S7F**). Importantly, metastasis-bearing mice receiving αCDH3-CAR T cells survived significantly longer than mice receiving un-transduced (UTD) or αCD19-CAR T cells (**Figure 7C)**.

**Fig. 7.**
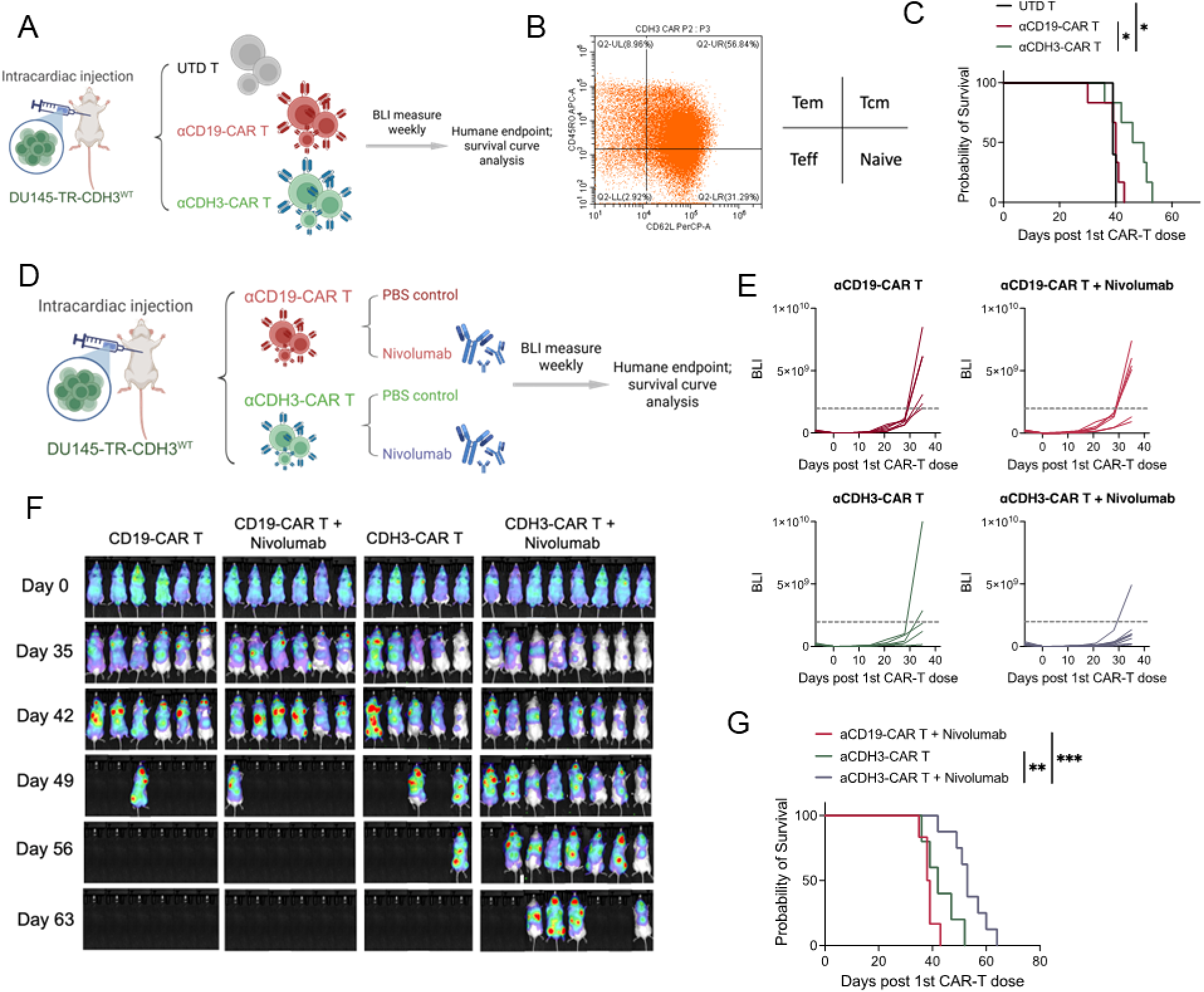
CDH3 targeted CAR T in vivo. (**A**) Schematic of the in vivo experimental design for metastatic prostate cancer models. DU145-TR are delivered via intracardiac injection, randomized into three groups, and treated with untransduced T (UTD T), αCD19-CAR T and) and αCDH3-CAR T cells. (**B**) Flow cytometric gating of infused CAR T cells, illustrating the proportions of key T-cell subsets (naïve, T effector, T effector memory, and T central memory). (**C**) Survival curves comparing UTD T cells, αCD19-CAR T, and αCDH3-CAR T treatment groups in metastatic prostate cancer model. (**D**) Schematic of the in vivo experimental design for combinational therapy. DU145-TR are delivered via intracardiac injection, randomized into four groups, and treated with αCD19-CAR T alone, αCD19-CAR T + nivolumab, αCDH3-CAR T alone, and αCDH3-CAR T + nivolumab. (**E**) Bioluminescence signaling intensity comparing four treatment group. (**F**) Bioluminescence imaging of the DU145-TR metastatic prostate cancer model, comparing the four treatment groups over several weeks. (**G**) Survival curves of αCDH3-CAR T alone, αCDH3-CAR T + nivolumab and αCD19-CAR T + nivolumab treatment groups. Log-rank (Mantel-Cox) test for (**C**) and (**G**). *P < 0.05, **P < 0.01, ***P < 0.001, ****P < 0.0001, ns, not significant.

Combining CAR T and ICB therapy is a highly pursued approach in clinical development. To test this strategy, we conducted another round of CAR-T therapy, alone or in combination with anti–PD1 antibody nivolumab (**Figure 7D, Figure S7E**). Metastasis burden tracked with bioluminescence imaging (BLI) and survival curves confirmed that αCDH3-CAR T therapy slowed metastasis progression, but more importantly, the combination of αCDH3-CAR T cells and nivolumab led to the lowest metastases and the longest survival (**Figure 7E-G, Figure S7H**).

In summary, CDH3-targeted CAR T cells demonstrated impressive antitumor activity against CDH3^+^ metastasis prostate cancer and elongated mouse survival. The efficacy was further enhanced by co-administration of anti-PD1, highlighting a promising combination approach.

## DISCUSSION

Our study establishes CDH3 as a pivotal biomarker and therapeutic target in basal-like double-negative prostate cancer. Starting with *Pten/Apc*–deficient mouse model, we showed that loss of these tumor suppressors drives metastatic prostate cancer marked by CDH3 overexpression and heightened WNT signaling. Corroborating human data indicate that CDH3 is significantly elevated in basal-like prostate cancer subtype, which are known for poor responses to androgen deprivation therapy. Mechanistically, we demonstrated that YAP1 and non-canonical WNT signals (WNT5A–ROR2) maintain CDH3 expression, linking CDH3 to key oncogenic pathways that promote invasiveness and treatment resistance. Therapeutically, we provide proof-of-concept that targeting CDH3 can effectively eliminate CDH3^+^ prostate cancer cells: a CDH3-targeted ADC delivered a cytotoxic payload to CDH3⁺ tumors with minimal impact on antigen-negative cells, and CDH3-specific CAR T cells mitigated CDH3^+^ metastases and improved survival in preclinical models. These results underscore the translational potential of CDH3–directed therapies for aggressive prostate cancer.

The findings reported here highlight a novel therapeutic opportunity for a molecularly defined subset of prostate cancer. Three key aspects of our work deserve emphasis. First, we applied a subtype-driven perspective by focusing on basal-like (double-negative) prostate cancer, which enabled the identification of CDH3 as a promising target. Basal-like tumors in prostate cancer have recently been recognized as an entity with worse prognosis and distinct biology compared to classical luminal androgen-driven tumors ^1,2^. A similar paradigm exists in other malignancies: for example, triple-negative breast cancers are often basal-like and respond poorly to standard treatments ^39 40^. Our approach demonstrates that honing in on the basal-like subtype – defined either by transcriptomic classifiers or by lack of lineage-specific markers – can unmask targetable vulnerabilities that might be overlooked in unstratified analyses. This strategy can be generalized to other heterogeneous cancers. Indeed, basal-like or “squamous” transcriptional subtypes have been described in pancreatic ductal adenocarcinoma and found to carry unique drug sensitivities ^41,42^. Likewise, muscle-invasive bladder cancers can be divided into basal and luminal subtypes with different chemotherapy responses ^43,44^, and head and neck squamous cell carcinomas comprise subgroups (including a basal-like cluster) with distinct clinical outcomes ^45,46^. By leveraging such subtype information, we can better tailor therapeutic discovery efforts. In the context of prostate cancer, our study validates the basal/double-negative classification as more than a prognostic label – it directly informed the successful targeting of CDH3.

Second, our integrative multi-omics approach proved powerful in pinpointing CDH3 as a crucial node in the aggressive tumor network. By combining proteomic (RPPA), transcriptomic (bulk RNA-seq), and single-cell analyses, we captured a comprehensive picture of how *Pten/Apc* loss reprograms the tumor. This revealed an interplay between CDH3 and the WNT/YAP pathways. Notably, our data show that CDH3 is not just a coincidental marker of basal-like cells but is actively maintained by the oncogenic signaling milieu (YAP and non-canonical WNT) prevalent in those cells. Such crosstalk exemplifies how a multimodal analysis can yield insights that single-technique studies might miss. The coordination between YAP, WNT5A/ROR2, and CDH3 highlights a regulatory circuitry driving tumor invasiveness and castration resistance. It aligns with prior knowledge that YAP can induce stem-like, therapy-resistant traits in prostate cancer^23^ and extends it by identifying CDH3 as a downstream effector of that program. More broadly, our study underscores that integrating data across different “omes” – proteome, transcriptome (bulk and single-cell) – can illuminate the key drivers of aggressive cancer phenotypes ^47^. Such integrative analyses will be increasingly vital as we attempt to map complex regulatory networks in heterogeneous tumors.

Third, from a translational standpoint, we demonstrate that CDH3 is an actionable target for therapy in prostate cancer. Basal-like prostate cancers typically do not benefit from hormonal therapies and currently have limited treatment options beyond chemotherapy. Our preclinical results suggest that CDH3-directed therapies could fill this gap by providing a way to selectively attack these tumors. In particular, ADCs against CDH3 could deliver cytotoxic agents directly to basal-like cancer cells while sparing normal tissues that lack CDH3. Supporting this, prior clinical investigations have explored CDH3 targeting in other cancers. For instance, an anti-CDH3 ADC (PCA062, by Novartis) was tested in a Phase I trial for solid tumors including TNBC and head & neck cancer; while that trial was terminated early due to limited efficacy, the treated patients were heavily pretreated end-stage cases and received few doses, which may have confounded outcomes ^48^. Another approach using a radiolabeled anti-CDH3 antibody (90Y-FF-21101) in advanced solid tumors showed good tumor uptake, though therapeutic effect was modest ^49^. These experiences suggest that CDH3 is a viable target, but optimal modalities and clinical contexts need to be identified. Our data support that improved ADC designs (e.g., using potent payloads like DM1 or MMAE with stable linkers) could yield better results, especially if applied in patients whose tumors highly express CDH3. Encouragingly, a new CDH3 ADC with a cleavable MMAE payload has shown promising activity in preclinical models of lung cancer and is entering clinical evaluation, indicating ongoing interest in this target.

We also show, for the first time to our knowledge, the feasibility of CAR T cells targeting CDH3. The CDH3-CAR T cells were very effective in our models, eradicating tumors and extending survival. Importantly, we did not observe overt toxicity in mice (noting that the models were immunodeficient). Safety will be a paramount concern for any CAR T cell targeting a tumor antigen that has some normal tissue expression. CDH3 is expressed in certain normal epithelia (e.g., skin, breast ducts, and stomach mucosa), but typically at lower levels than in carcinomas. Early clinical data hint that on-target toxicities of CDH3 therapies (ADC or radioimmunotherapy) are manageable ^48 49^. Nonetheless, careful design of dosing and perhaps transient CAR T control strategies (such as a suicide switch) may be prudent for translating CDH3-CAR T cells. The fact that our CAR T cells showed enhanced efficacy when combined with PD-1 blockade is noteworthy. Checkpoint inhibitors could be used adjunctively to boost CAR T function in solid tumors, as we demonstrated. This combined approach might surmount some of the immunosuppressive barriers in the tumor microenvironment, leading to more durable remissions.

One limitation of our study is the incomplete assessment of potential toxicities and the long-term effects of CDH3 targeting. While our preclinical models are promising, clinical translation will require ensuring that normal tissues with low CDH3 expression (e.g., skin basal cells or hair follicles) are not unduly harmed. Early trials of CDH3-targeted agents reported mostly mild toxicities and no dose-limiting damage to normal epithelia ^48 49^, which is encouraging. Still, further investigation in relevant models (including primates, if possible) would be valuable to fully de-risk this approach. Another point is that we focused on CDH3’s role in cancer cells; however, CDH3 could also influence the tumor microenvironment (for example, by mediating cell–cell interactions or affecting immune infiltration).

Looking ahead, our findings open several avenues. On the mechanistic side, further dissection of the YAP–WNT5A–CDH3 loop in basal-like cancers could reveal additional targets. The fact that CDH3 is downstream of YAP and WNT signaling makes it a convenient point of therapeutic attack, but other nodes in the loop (e.g., FOSL1 or ROR2) might also be targetable to synergize with CDH3 therapies. On the therapeutic side, our results strongly motivate clinical trials of CDH3–directed therapies in biomarker-selected patients. Basal-like or double-negative prostate cancers could be identified via gene expression profiling or IHC (for example, low AR and high cytokeratin 5/6 or CDH3 itself). Patients with metastatic castration-resistant disease falling into this category might particularly benefit from a CDH3-ADC or CAR T cell therapy, as conventional options are scarce. Additionally, combining modalities – as illustrated by CAR T plus PD-1 blockade – may enhance outcomes and should be explored.

In conclusion, this work provides a blueprint for translating a biological insight (CDH3 upregulation in basal-like prostate cancer) into therapeutic innovation. It underscores the value of tumor subtyping and multi-omics integration in discovering actionable cancer targets. By demonstrating that CDH3 can be effectively targeted with advanced immunotherapeutics, we lay the groundwork for new treatment strategies aimed at improving the prognosis of patients with aggressive, treatment-refractory prostate cancer.

## MATERIALS AND METHODS

### Study design

This study aimed to define and validate CDH3 as a therapeutic target in basal-like prostate cancer and triple-negative breast cancer (TNBC). The study consisted of three major components. First, we developed a genetically engineered mouse model (Pb^Cre^Pten^L/L^Apc^L/L^ROSA^mT/mG^) to explore the cooperative role of Pten and Apc loss in driving aggressive tumor phenotypes, with CDH3 upregulation characterized by transcriptomic and proteomic analyses. Second, we validated the association of CDH3 with basal-like features in a castration-resistant prostate cancer (CRPC) cohort and confirmed its overexpression in public prostate cancer transcriptomic datasets using PAM50 subtyping. Single-cell RNA sequencing was further employed to establish the clinical relevance and cellular specificity of CDH3 expression. Third, we evaluated CDH3 as a therapeutic target by testing the efficacy and specificity of a CDH3-targeted antibody-drug conjugate (ADC) and CDH3-specific chimeric antigen receptor (CAR) T cells. In vitro assays measured antigen-dependent internalization, cytotoxicity, and T cell activation, while in vivo experiments assessed tumor control and survival benefit in metastatic prostate and breast cancer models treated with ADC or CAR T cells, with or without PD-1 blockade. All in vitro experiments were performed using biological triplicates and repeated at least twice, with statistical details provided in the figure legends. In vivo experiments were conducted using immunocompromised mice (nude or NOD-SCID) aged 8 to 12 weeks. Tumor-bearing mice were randomized into treatment and control groups. Metastatic models were generated via intracardiac injection of luciferase-labeled prostate or breast cancer cells, followed by treatment with ADC or CAR T cells. End points included bioluminescence imaging, tumor burden, and survival. No data were excluded from analysis. Group sizes (typically n = 5–10 mice) were selected based on prior studies and power analysis.

### Mice

PB-Cre4, PtenL/L, Smad4L/L, p53L/L, and ROSAmT/mG alleles have been previously described. The ApcL/L mice (Jackson Lab) were purchased for cross to generate Pb^Cre^Pten^L/L^Apc^L/L^ROSA^mT/mG^ mice. All GEM mouse models were backcrossed to a C57BL/6 background for at least four generations. NOD-SCID mice (IMSR_JAX:001303) were purchased from the Jackson Laboratory. Nude mice (RRID: IMSR_TAC:ncrnu) were purchased from Taconics. All animal work performed in this study was approved by the Institutional Animal Care and Use Committee (IACUC) at University of Notre Dame (protocol number 20-03-5930). All animals were maintained under pathogen-free conditions and cared for in accordance with the International Association for Assessment and Accreditation of Laboratory Animal Care policies and certification.

### Reverse phase protein array (RPPA)

RPPA was conducted by the RPPA Core Facility at MD Anderson Cancer Center following the standard protocols, the details of which can be found on the core facility website (www.mdanderson.org/research/research-resources/core-facilities/functional-proteomics-rppa-core.html). The dataset returned from the facility included raw data in log2 value and normalized data in linear value.

### Microarray and analysis

RNA was purified from whole tumor lysate or sorted GFP+ tumor cells, as described in Figure S1C, using RNeasy Mini Kit (Qiagen, Germantown, MD, USA). Microarray (Affymetrix Mouse Genome 430 2.0 Array) was conducted by the Sequencing and Microarray Facility at MD Anderson Cancer Center following the facility’s standard procedure. The data were analyzed in RStudio, including oligo package and arrayQualityMetrics package used for quality control and calibration, ArrayExpress package and mouse4302.db package used for probe annotation, and package was used for differential expression analysis. p value of 0.05 or 0.01 by unpaired t-test was used as threshold of significantly upregulated or downregulated genes by from different genotypes. For pathway analysis, the gene lists were imported to RStudio with clusterProfiler package used for KEGG, Wiki and Hallmark 50 pathway analysis.

### CDH3 targeting antibody and ADC

The anti-CDH3 antibody TSP44d (human IgG), targeting the extracellular cadherin domain 1 (ECD1) of human CDH3, was developed and gifted by Perseus Proteomics Inc. The antibody-drug conjugate (ADC) was synthesized by conjugating TSP44d to the microtubule inhibitor DM1 via a non-cleavable SMCC linker (TSP44d–SMCC–DM1), also by Perseus Proteomics Inc. The resulting ADC had a predominant drug-to-antibody ratio (DAR) of 2–4, measured with mass spectrometry.

### Cell lines and genetic modification

The DU145, 22RV1, LHMK, PC3, LNCaP and Jurkat cell lines were cultured in RPMI 1640 Medium (Gibco) supplemented with 10% heat-inactivated fetal bovine serum (FBS, Gibco) and 1% penicillin/streptomycin mix (P/S, Gibco). CDH3 and ROR2 were knocked out by lentivirus transduction using CRISPR/Cas9 All-in-One Lentivector set (Applied Biological Materials).

### HE and IHC

Formalin-fixed, paraffin-embedded (FFPE) tissue sections (5 µm) were deparaffinized in xylene, rehydrated through graded ethanol, and subjected to either hematoxylin and eosin (H&E) staining or immunohistochemistry (IHC). For H&E, sections were stained with Mayer’s hematoxylin for 8 min, rinsed in tap water, counterstained with Eosin Y (VWR, 1% alcoholic) for 10–15 min, dehydrated, cleared in xylene, and mounted with Permount (Fisher Scientific). For IHC, antigen retrieval was performed in citrate buffer (pH 6.0) using a pressure cooker. After blocking with 5% goat serum, sections were incubated overnight at 4 °C with anti-CDH3 antibody (R&D Systems, AF761), followed by HRP-conjugated secondary antibody (ImmPRESS, Vector Labs) and DAB substrate (Vector Labs). Nuclei were counterstained with hematoxylin, and slides were dehydrated and mounted.

### Flow cytometry

For all the samples, cells were preincubated with 1:200 human Fc block (brand) and mouse Fc block (band) in fluorescent-activated cell sorting (FACS) buffer consisting of PBS with 2% fetal bovine serum (FBS, brand) and 2mM EDTA. Cells were then stained with antibodies against surface markers in the FACS buffer at 4°C for 30 min. In all experiments, cells were also labeled with 1:1000 DAPI (brand) in FACS buffer at 4C for 10 min and washed and resuspended in FACS buffer before acquisition on Beckman CytoFLEX.

### TSS44d Ab / TSS44d-DM1 labelling and internalization study

Naked anti-CDH3 Ab (TSS44d) or ADC were labelled with pHordoTM (Thermo) that excited the red fluorescent under low pH. pHordo labeled antibody or ADC were diluted at the different concentrations in cell culture medium and incubate with DU145-TR-CDH3^WT^ or DU145-TR-CDH3^KO^ cells for 48 hours. Internalization of antibody or ADC were visualized by fluorescent microscope (Leica) and quantified by flow cytometry (Beckman CytoFLEX).

### In-vitro treatment of ADC and proliferation assay

Cells were seeded into appropriate culture plates, and antibody-drug conjugates (ADC) were prepared at various concentrations by dilution in the respective culture medium for each cell type. The cells were then incubated with the ADC-containing medium for 72 hours at standard culture conditions. After incubation, the ADC medium was aspirated, and cells were incubated with a medium containing 1.5 mg/mL MTT reagent for 4 hours. Following the MTT incubation, the medium was removed, and the resulting purple formazan crystals were dissolved by adding dimethyl sulfoxide (DMSO). The absorbance of the dissolved formazan was measured at 570 nm with a reference wavelength of 630 nm using a plate reader (company, model). The half-maximal inhibitory concentration (IC50) values were determined using GraphPad Prism software v10.

### Design of the anti-CDH3 CAR molecule

The anti-CDH3 chimeric antigen receptor (CAR) was designed using the single-chain variable fragment (scFv) derived from the TSS44d Ab used in the ADC. The DNA sequence encoding the scFv was codon-optimized for human expression and synthesized by Genscript. The synthesized DNA fragment was subsequently cloned into the backbone of pSLCAR-CD19-BBz (Addgene #135992), replacing the original anti-CD19 scFv. This plasmid contains a CD8α hinge and transmembrane domain, 4-1BB costimulatory domain, and CD3ζ signaling domain. Standard molecular cloning techniques, including restriction enzyme digestion and ligation, were used to generate the final CAR construct, which was verified by Sanger sequencing.

### CAR-Jurkat cell transduction and activation

Lentiviral particles encoding either the anti-CDH3 CAR or the anti-CD19 CAR (pSLCAR-CD19-BBz) were produced and used to transduce Jurkat T cells. Transduction efficiency was assessed 72 hours post-infection by measuring GFP expression, which is encoded on the plasmid backbone, using flow cytometry. To evaluate CAR-mediated activation, transduced Jurkat cells were co-cultured with either DU145-CDH3^WT^ cells or DU145-CDH3^KO^ cells at a 5:1 (Jurkat to cancer cell) ratio for 24 hours. Surface expression of activation markers CD69 and CD25 on Jurkat cells was then assessed by flow cytometry. Activation was quantified based on the percentage of CD69+ and CD25+ cells within the GFP+ (CAR-expressing) Jurkat population.

### Human CAR-T cell Generation

Frozen human peripheral blood mononuclear cells (PBMCs) were purchased from STEMCELL. Upon thawing, PBMCs were resuspended in T cell culture medium (TCM), composed of X-VIVO 15 Medium (Lonza #BE02-060F) supplemented with human IL-7 (R&D Systems #P13232), IL-15 (R&D Systems #P40933), and IL-21 (Novoprotein #GMP-CC45), and rested at 37 °C for 1-2 hours. To enrich T cells, the thawed PBMCs were allowed to settle by gravity, and the non-adherent supernatant (enriched in T lymphocytes) was collected for further processing. T cells were stimulated for 48-72 hours using anti-hCD3 and anti-hCD28 antibody-coated immunobeads at a 1:1 bead-to-cell ratio in TCM. Activated T cells were transduced with lentiviral particles encoding either αCDH3-CAR or αCD19-CAR in the presence of LentiBOOST (Mayflower Bioscience, SB-P-LV-101-03), which was added according to the manufacturer’s instructions to enhance transduction efficiency. Lentiviral transduction was performed at room temperature with centrifugation at 1000×g for 1.5 hours, followed by incubation overnight. The culture medium was replaced with fresh TCM the next day. Immunobeads were removed for 6-7 days post-transduction. Transduced T cells were then expanded in TCM, with medium replenished and cell density adjusted to 0.5-2 × 10⁶ cells/mL every 2-3 days. Expanded CAR-T cells were rested and used for downstream assays.

### In-vitro CAR T activation experiments

To evaluate antigen-specific activation, human CAR-T cells were co-cultured with either DU145-CDH3^WT^ cells or DU145-CDH3^KO^ cells at an effector-to-target (E:T) ratio of 5:1 in TCM without cytokine supplementation. After 24 hours of co-culture at 37 °C, cells were harvested and stained for surface markers to identify CAR-T cells, followed by fixation and permeabilization for intracellular staining. Intracellular expression of Granzyme B and Perforin was assessed using flow cytometry to quantify the activation and cytolytic potential of CAR-T cells.

### In-vitro CAR T killing experiment

To assess the tumor-killing ability of CAR-T cells, in vitro co-culture experiments were performed with DU145-CDH3^WT^ or DU145-CDH3^KO^ target cells. Tumor cells were seeded in 96-well plates and allowed to adhere overnight. Human CAR-T cells were then added at varying effector-to-target (E:T) ratios, and co-cultured for 72 hours in TCM without exogenous cytokines. Following the incubation period, non-adherent cells (including T cells) were carefully removed by gentle washing. Tumor cell viability was then assessed using the MTT assay, as described previously in the ADC treatment section.

### Metastasis mice models

Groups of five to seven nude or NOD-SCID mice aged between 8-12 weeks were injected intracardiacally with DU145 cells in 50uL of PBS. The DU145 cells were labeled with a triple reporter (TR) of the HSV1-tk-GFP-luciferase fusion protein^33^. To monitor the tumor burden, 100uL of 15mg/mL D-luciferin in PBS were intravenously injected in the mice and the bioluminescent intensity (BLI) was measured by SPECTRAL Ami HT in vivo imaging system.

### In vivo treatment of ADC on mouse metastatic cancer model

For ADC treatment on male mice, 2 × 105 DU145 cells were intra-cardiac injected into male nude mice on D0. ADC at 2.5mg/kg or 5mg/kg or equivalent volume of PBS were intravenously injected twice a week from D12. Tumor burden was monitored through the measuring of BLI signals.

### CAR-T therapy on mouse metastatic cancer model

2 × 105 DU145 cells were intra-cardiac injected on D0. All of the mice were preconditioned with cyclophosphamide (Cy) at 100mg/kg by intraperitoneal injection on D7. Mice received 3 × 106 αCDH3-CAR or αCD19-CAR T cells intravenously on D8, D9 and D10 with human IL-2 (Pepro Tech)) support at 30,000 IU per injection on D8, D9, D10, D12, D14. In the anti-PD-1 experiment groups, groups of mice were also treated with human anti-PD-1 Ab (Nivolumab, Leinco Technologies) at 200ug per injection at D8, D12, D14, D17. Tumor burden was monitored through the measuring of BLI signals.

### Public prostate cancer patient transcriptomic analysis

RNA-seq data of Tang et al. were downloaded from GSE199190, and we used the same groups of samples from the literature, with code available from the literature with some modification. RNA-seq gene expression data and clinical information of TCGA PRCA and the Fred Hutchinson Cancer Research Center (FHCRC) were downloaded from cBioportal. The analysis of the RNA-seq data followed the pipeline described by Law et al. The following four prostate datasets of microarray are downloaded from Gene Expression Omnibus (GEO): Mayo Clinic (GSE46691), Cleveland Clinic (GSE62267), Thomas Jefferson University (GSE72291), John Hopkins Hospital (GSE79958). The data processing was similar as mentioned above for mouse microarray data, except for annotation of human transcriptomic probes is by package HuEx-1_0-st-v2.

### PAM50 clustering

PAM50 subtyping was performed using the original algorithm by Parker et al., with gene expression data median-centered per cohort. The normal-like and HER2-enriched subtypes were excluded due to limited normal tissue and lack of ERBB2 amplification in prostate cancer, respectively. Samples were classified as luminal A, luminal B, or basal based on the highest correlation.

### Public prostate cancer patient single-cell transcriptomic analysis

Single cell RNA-seq analysis of public human prostate cancer datasets (GSE176031, GSE 185344, GSE221603) was performed by using ‘Seurat’ R package, including data quality control, filtering and normalization. Cells are annotated into epithelial cell, immune cell and stroma cell groups by ‘scType’ and ‘SingleR’ package with fine tune based on literature mining.

### Statistical analysis

All statistical analyses (as specified in the figure legends) were performed in Graphpad Prism v10. Data analyses were conducted using unpaired Mann-Whitney test to compare two groups or using Kruskal-Wallis when analyzing more than two sets of data.

## ACKNOWLEDGMENT

We would like to thank the Lu lab members for helpful comments and suggestions during this work. We are grateful for the support from various core facilities at Notre Dame, especially Freimann Life Science Center and Integrated Imaging Facility.

## AUTHOR CONTRIBUTIONS

G. Liu: conceptualization, investigation, methodology, data curation, formal analysis, validation, visualization, project administration, writing – original draft, writing – review & editing. S. Wang, L. Doung, Z. Zeng: investigation, methodology, data curation, formal analysis; J. Yang, Y. Zhao, S. Rahmy, S. Feng, A. Arce, L. Jia: investigation; J. Wan, L. Cheng, Xuemin Lu: supervision, funding acquisition; Xin Lu: conceptualization, investigation, resources, supervision, funding acquisition, writing – original draft, writing – review & editing.

