## Supplemental Figures for "CDH3 as a Novel Therapeutic Target in Basal-like Double-Negative Prostate Cancer"

**Figure S1. Cdh3 upregulation and WNT pathway dysregulation in the Pten/Apc prostate specific knockout mouse.**

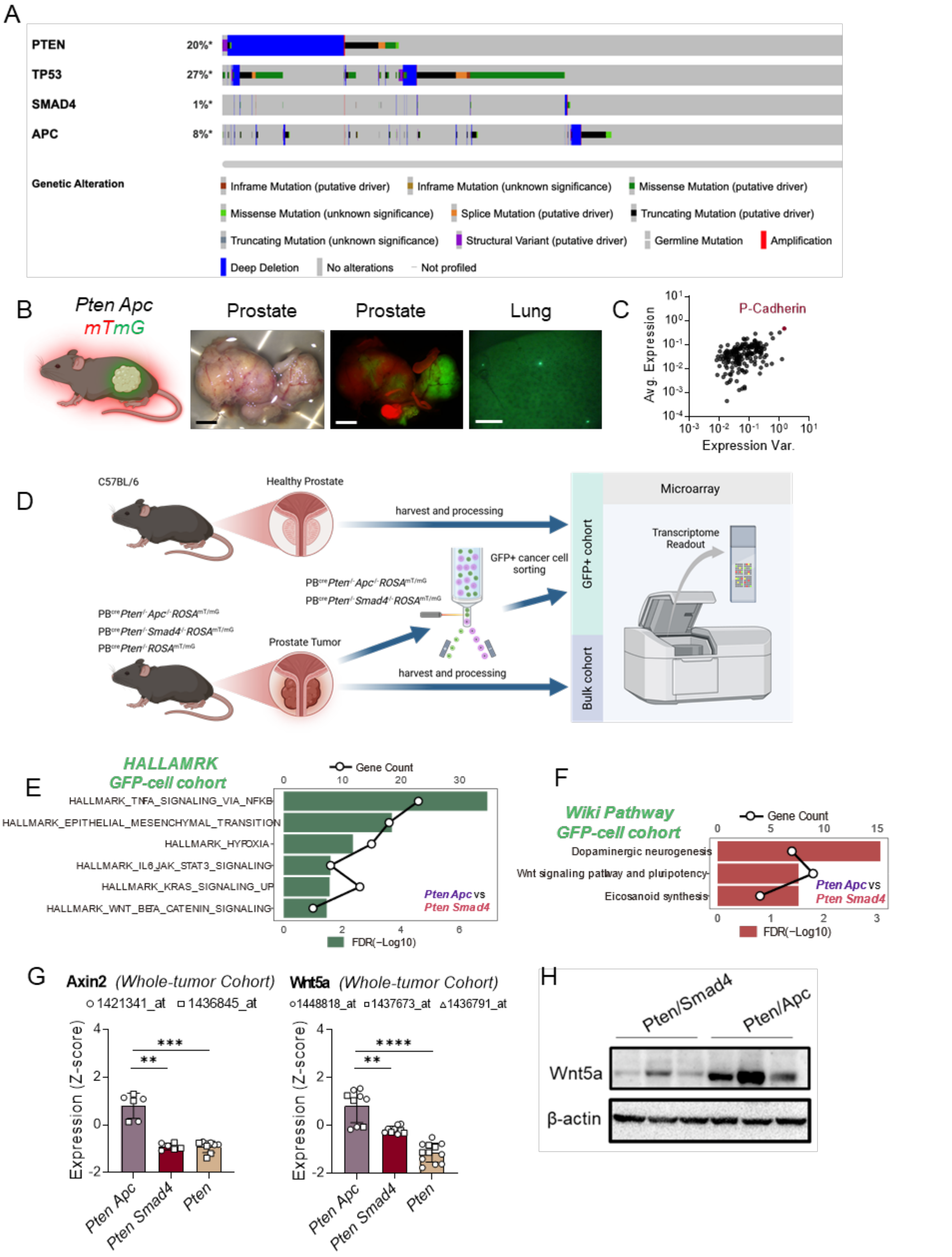

(A) Genomic overview of key tumor suppressor gene alterations-PTEN, TP53, SMAD4, and APC-identified in prostate cancer datasets from cBioPortal.

(B) Representative prostate and lung tissues from $PB^{Cre}Pten^{L/L}Apc^{L/L}ROSA^{mT/mG}$ mice at approximately 4 months of age. Cre-expressing tumor cells appear green, while non-tumor cells remain red (tomato). Scale bar: 5 mm for prostate; 1 mm for lung.

(C) Scatter plot showing the average protein expression versus expression variability across 281 proteins in all samples. P-Cadherin, the protein with the highest differential expression, is highlighted in red.

(D) Schematic of the sample collection strategy for whole-tumor (bulk) and GFP+ cohorts: prostate tumors were harvested from corresponded genotypes and control animals, dissociated, and subjected to fluorescence-activated cell sorting (FACS) to separate GFP-expressing cancer cells from the bulk population. RNA was then extracted and analyzed via microarray, generating data for downstream pathway enrichment.

(E) Wiki Pathways analysis of the GFP+ tumor cell transcriptomes (*Pten Apc* vs. *Pten Smad4*) using genes significantly upregulated (log2FC > 2.5, adj. p-value < 0.05), highlighting significant enrichment for dopaminergic neurogenesis, Wnt signaling/pluripotency, and eicosanoid synthesis pathways (bars show FDR-adjusted p-values; circles indicate the number of differentially regulated genes).

(F) Hallmark gene set enrichment results for the GFP+ tumor cell transcriptomes (*Pten Apc* vs. *Pten Smad4*) using genes significantly upregulated (log2FC > 2.5, adj. p-value < 0.05), , indicating activation of multiple oncogenic programs-including TNFα signaling, epithelial-mesenchymal transition (EMT), hypoxia, IL6-JAK-STAT3, and notably Wnt-β-catenin signaling.

(G) Corresponding Axin2 and Wnt5a expression analysis in the whole-tumor cohort across different genotypes.

(H) Western blot validation of Wnt5a upregulation in *Pten Apc* tumor lysates compared to *Pten Smad4*.

Error bars represent SEM. Mann-Whiney test (black asterisks) for (**G**). *P < 0.05, **P < 0.01, ***P < 0.001, ****P < 0.0001.

**Figure S2. CDH3 upregulation in treatment resistant basal-like prostate cancer**

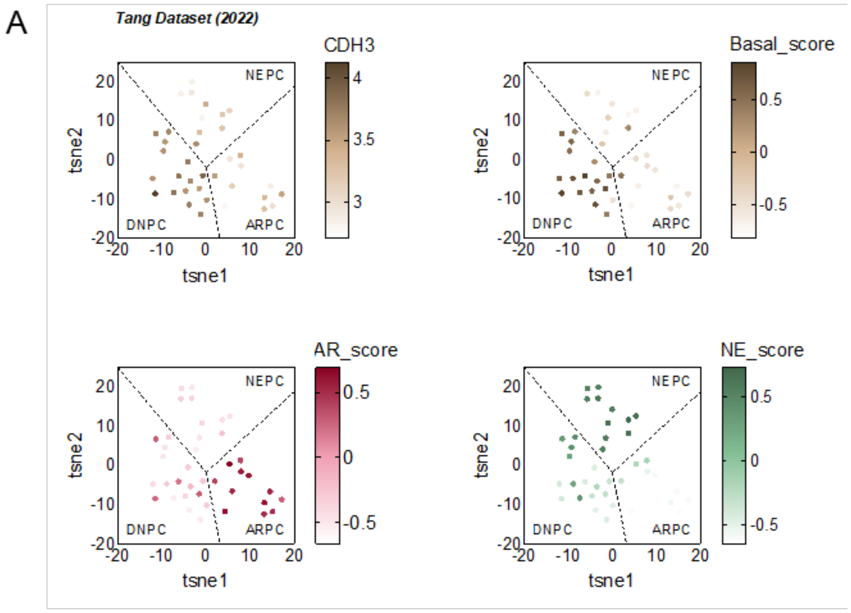


(A) t-SNE plots from the Tang et al. (2022) dataset. Each panel is overlaid with different signature scores (AR, NE, SCL, luminal, basal) or CDH3 expression. Darker colors indicate higher scores/expression.

**Figure S3. CDH3 has Preferential Association with Non-canonical WNT signaling in basal cells.**

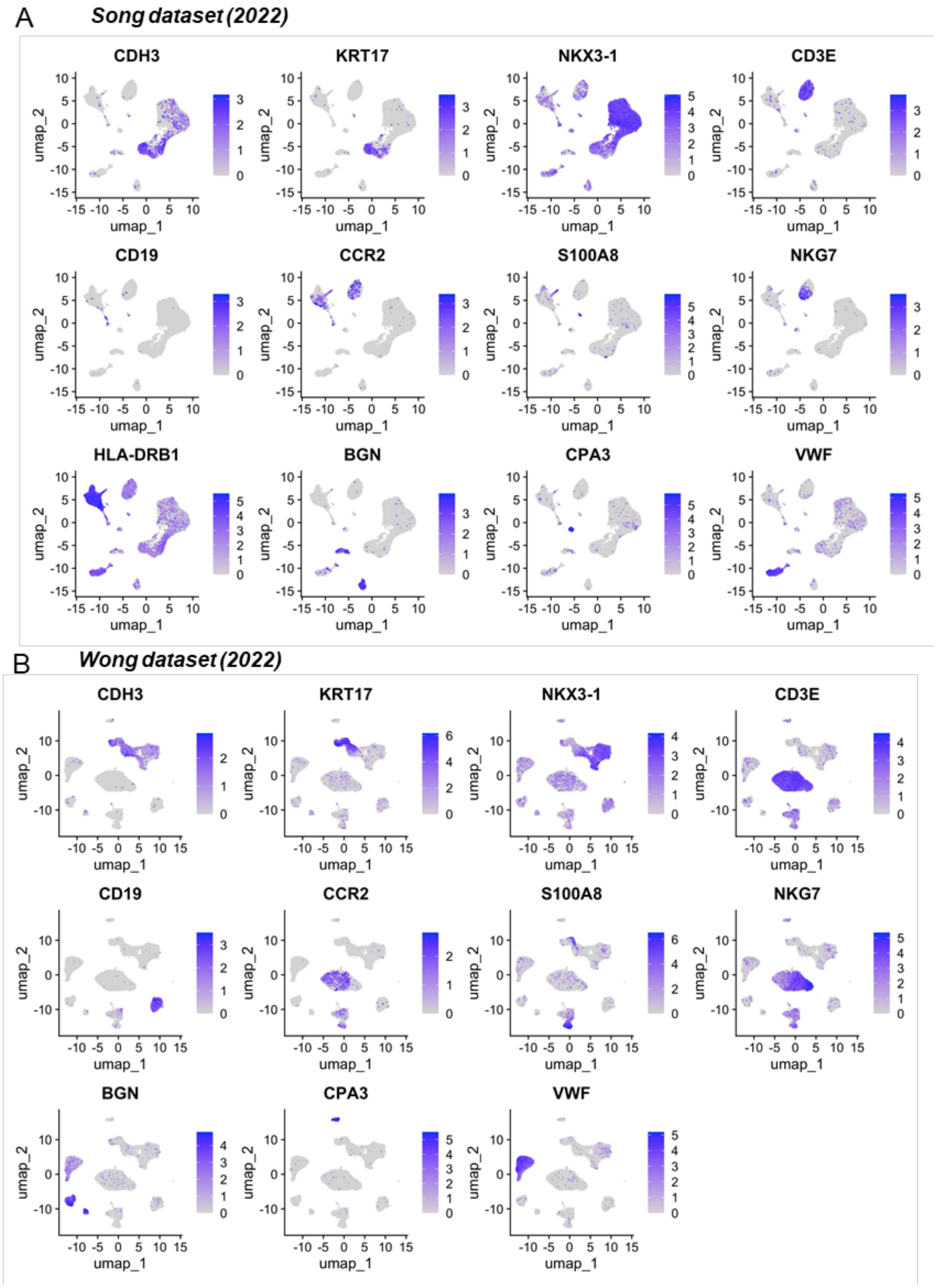

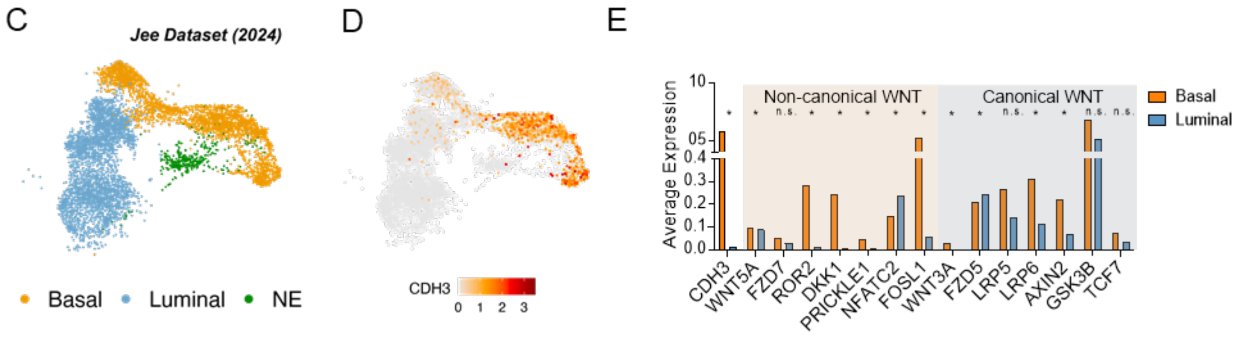


(A) UMAP plots of single-cell RNA-seq data from prostate cancer samples in the Song et al. (2022) dataset, with each panel showing the expression level (purple gradient) of a signature gene. Markers include CDH3, KRT17(basal epithelial cells), and NKX3-1 (luminal epithelial cells), CD3E (T cells), CD19 (B cells), CCR2/S100A8 (myeloid), NKG7 (NK cells), HLA-DRB1 (antigen-presenting cells), BGN (fibroblasts), CPA3 (mast cells), and VWF (endothelial). Higher intensity (darker purple) indicates stronger expression, facilitating cell-type annotation within the tumor microenvironment.

(B) Similar UMAP plots for Wong et al. (2022) prostate cancer dataset.

(C) Single-cell UMAP from the Jee et al. (2024) dataset showing prostate cancer cells grouped into basal epithelial, luminal epithelial, neuroendocrine (NE) population.

(D) CDH3 expression in feature plot (heatmap scale).

(E) Comparison of average expression of canonical vs. non-canonical WNT pathway components in basal vs. luminal cells.

**Figure S4. CDH3 regulation through YAP / ROR2**

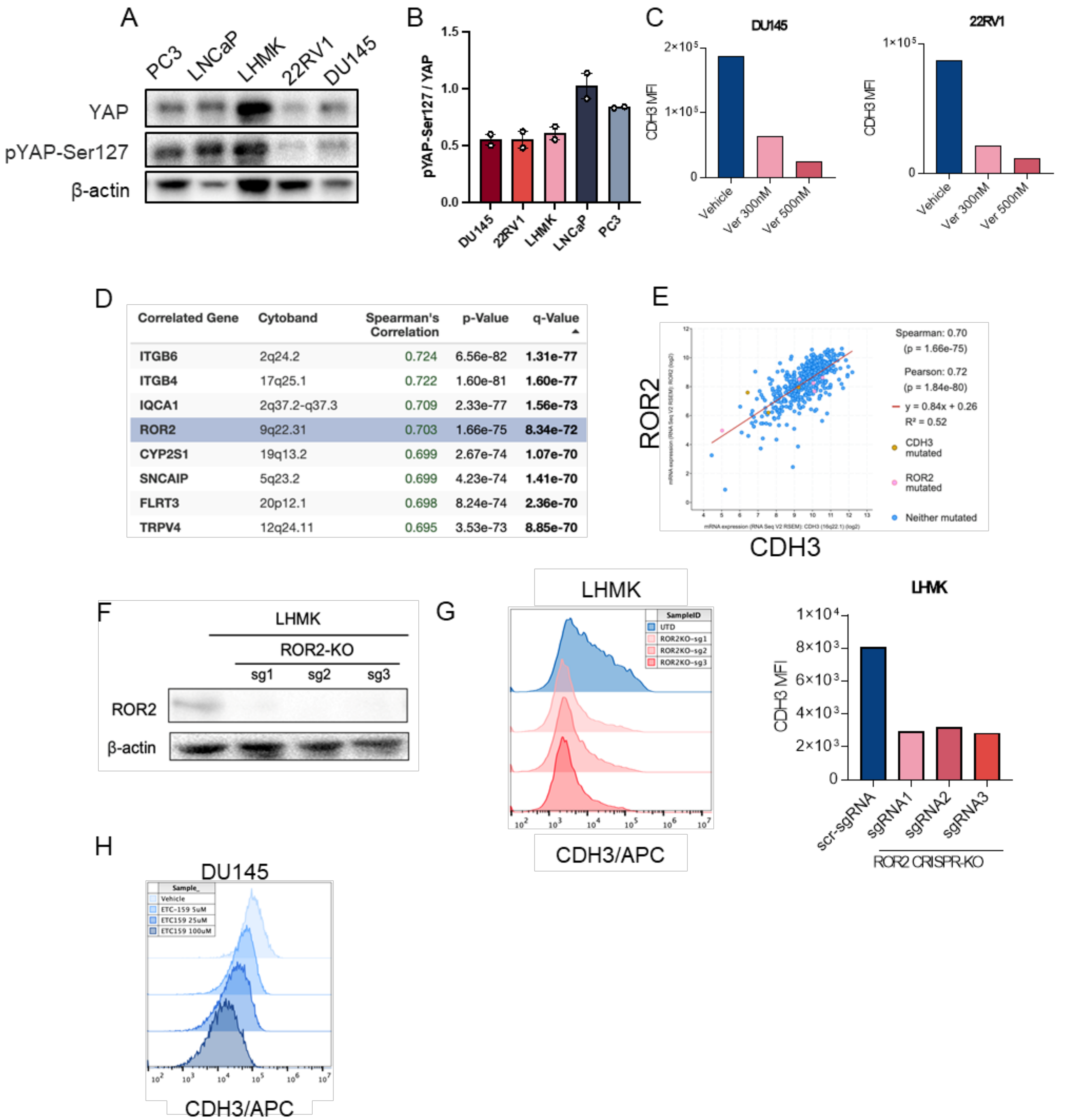


| Correlated Gene | Cytoband | Spearman's Correlation | p-Value | q-Value |
|---|---|---|---|---|
| ITGB6 | 2q24.2 | 0.724 | 6.56e-82 | 1.31e-77 |
| ITGB4 | 17q25.1 | 0.722 | 1.60e-81 | 1.60e-77 |
| IQCA1 | 2q37.2-q37.3 | 0.709 | 2.33e-77 | 1.56e-73 |
| ROR2 | 9q22.31 | 0.703 | 1.66e-75 | 8.34e-72 |
| CYP2S1 | 19q13.2 | 0.699 | 2.67e-74 | 1.07e-70 |
| SNCAIP | 5q23.2 | 0.699 | 4.23e-74 | 1.41e-70 |
| FLRT3 | 20p12.1 | 0.698 | 8.24e-74 | 2.36e-70 |
| TRPV4 | 12q24.11 | 0.695 | 3.53e-73 | 8.85e-70 |

(A) Western blot for total YAP and phosphorylated YAP (pYAP-Ser127) in a panel of prostate cancer cell lines (PC3, LNCaP, LHMK, 22RV1, DU145) with β-actin as a loading control.

(B) Western-blot based quantification of the pYAP-Ser127/YAP ratio in each cell line, highlighting differences in YAP activation status.

(C) Flow cytometry analysis (MFI) of CDH3 following 72 h verteporfin (Ver) treatment in DU145 and 22RV1 cells. Increasing concentrations of verteporfin (300 nM, 500 nM) lead to a dose-dependent reduction in CDH3 expression.

(D) Table from the cBioportal analysis of TCGA prostate cancer (PRCA), listing the top genes correlated with CDH3 expression. ROR2 appears among the highest correlated genes.

(E) Scatter plot comparing CDH3 and ROR2 transcript levels in the same dataset, indicating a strong positive correlation (Spearman's r = 0.70, Pearson's r = 0.72). Points are colored by whether CDH3 or ROR2 is mutated, with the best-fit regression line shown in red.

(F) Western blot confirming ROR2 knockout in LHMK cells (sgRNA1–3 vs. control), with β-actin as a loading control.

(G) Flow cytometry histogram and corresponding bar plot of CDH3 MFI in LHMK cells following ROR2 CRISPR-KO.

(H) Dose-dependent decline in CDH3 expression in DU145 cells treated with increasing concentrations of ETC159 (5-100 μM), shown by flow cytometry histograms.

**Figure S5. Effect of CDH-targeted ADC on prostate cancer cells.**

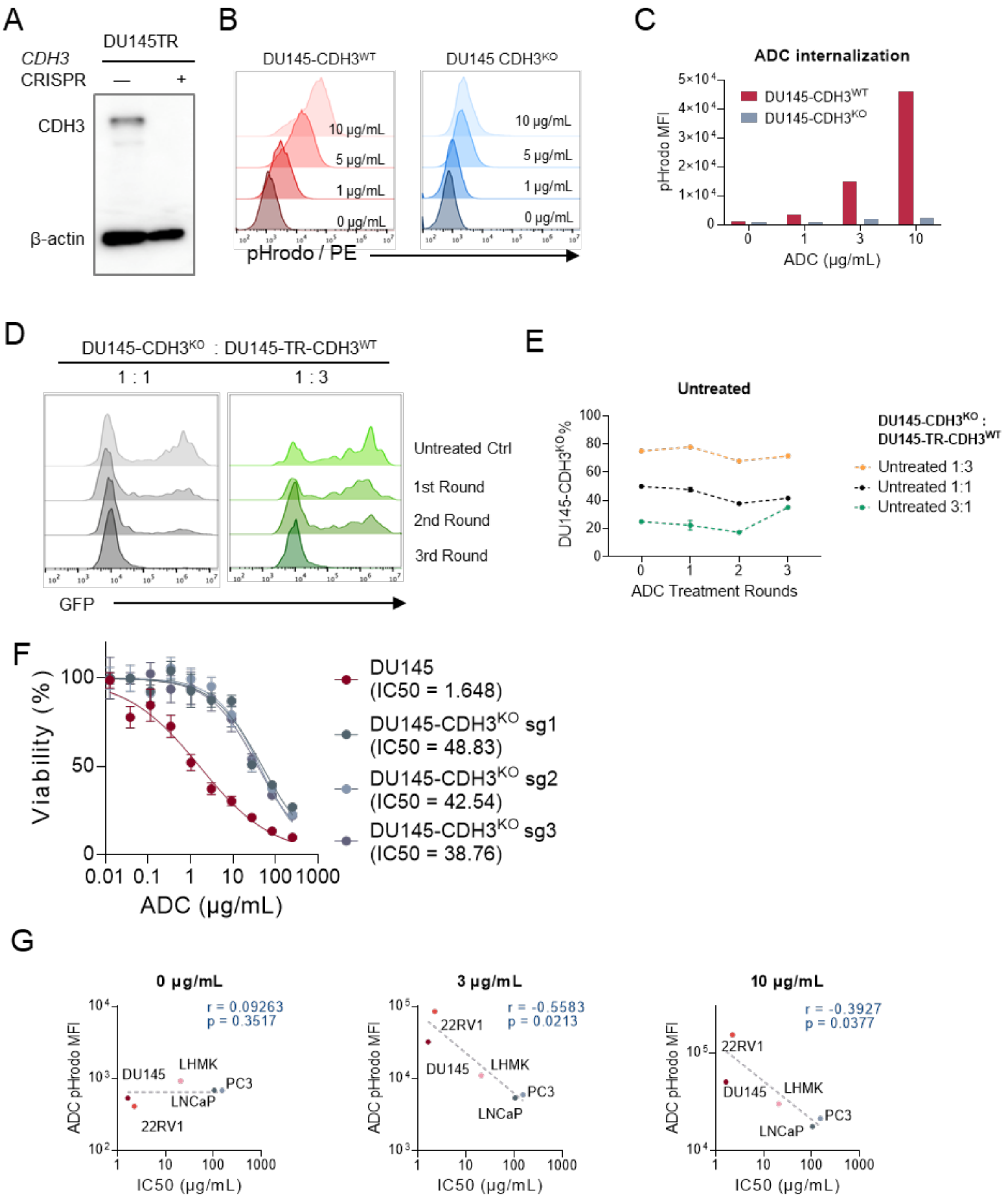

(A) Western blot showing CDH3 is knocked out in the DU145 cell line

(B) Flow cytometric histograms depicting pHrodo-labeled ADC uptake in DU145-CDH3$^{WT}$ vs. DU145-CDH3$^{KO}$ cells, measured at increasing ADC concentrations (0-10 μg/mL).

(C) Quantification of the MFI of pHrodo-labeled ADC in DU145-CDH3$^{WT}$ vs. DU145-CDH3$^{KO}$ cells.

(D) Flow cytometric histograms tracking GFP+ (DU145-CDH3$^{WT}$) and GFP- (DU145-CDH3$^{KO}$) populations at baseline (untreated) and after each ADC treatment round, presented for 1:1 and 1:3 initial ratios.

(E) Plot depicting changes in CDH3$^{KO}$ cell percentage over three rounds of culture without ADC (untreated control) for 1:3, 1:1 and 3:1 initial ratios.

(F) Dose–response cytotoxicity curves for CDH3$^{WT}$ and CDH3$^{KO}$ DU145 clones (sg1, sg2, sg3) treated with an anti-CDH3 ADC at increasing concentrations (0.01-300 μg/mL).

(G) Scatter plots illustrating the relationship between ADC internalization (measured by pHrodo mean fluorescence intensity, MFI) and cytotoxicity ($IC_{50}$) across various prostate (22RV1, DU145, LHMK, PC3, LNCaP) cancer cell lines. Each panel corresponds to a different ADC concentration (0, 3, or 10 μg/mL).

**Figure S6. Effect of CDH3-targeted CAR T cells on prostate cancer cells.**

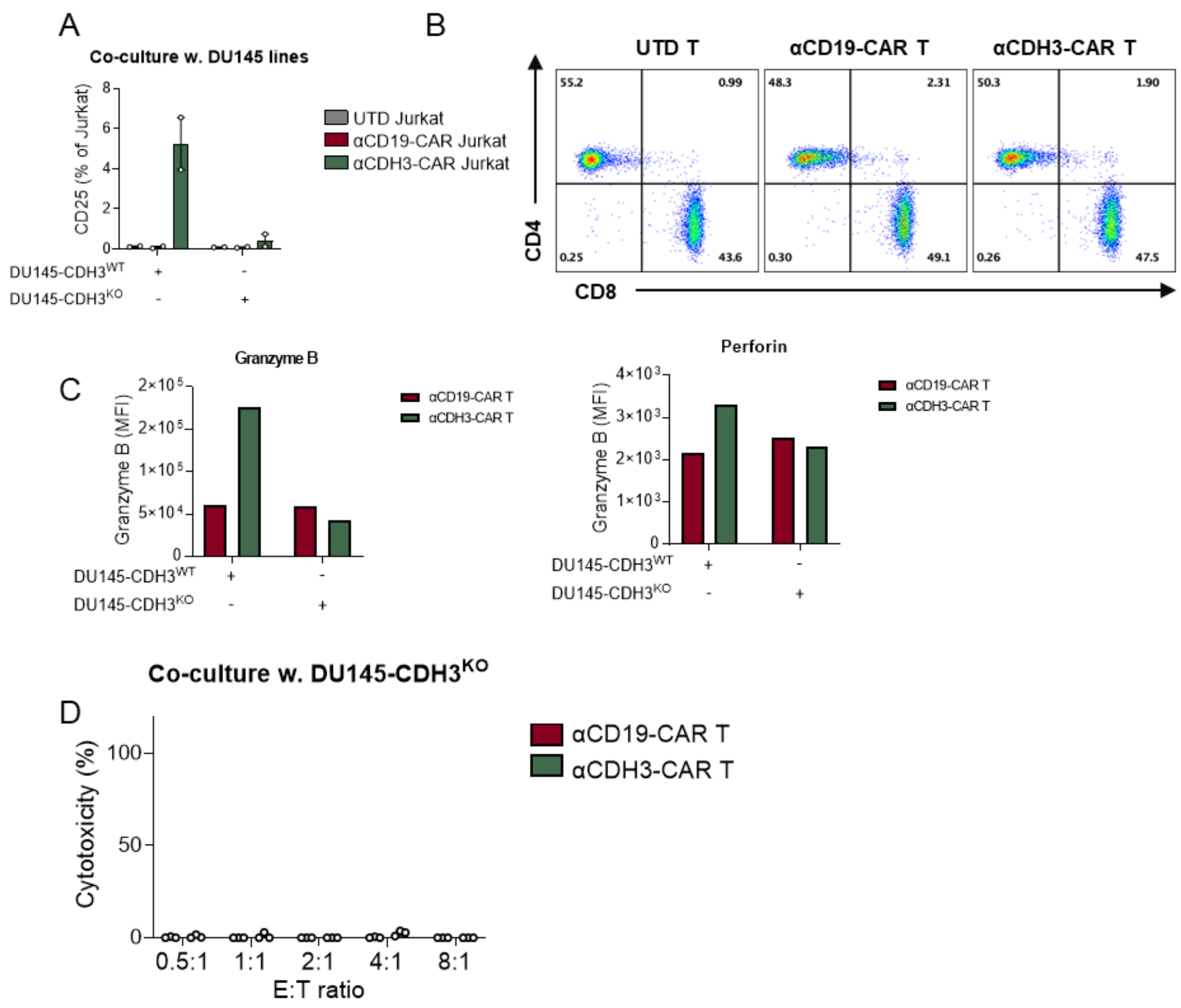

(A) Flow cytometry–based quantification of Jurkat cell activation (CD25 expression) following coculture with DU145-CDH3$^{WT}$ or DU145-CDH3$^{KO}$ cells, comparing the αCDH3-CAR with untransduced (UTD) and CD19-CAR control.

(B) Representative flow cytometry dot plots showing CD4 vs. CD8 portions in UTD T cells, αCD19-CAR T cells, or αCDH3-CAR T cells

(C) Flow cytometry analysis (NFI) of Granzyme B and perforin expression in αCD19-CAR T cells vs. αCDH3-CAR T cells upon co-culture with DU145-CDH3$^{WT}$ or DU145-CDH3$^{KO}$ cells

(D) Cytotoxicity assays against DU145-CDH3$^{KO}$ cells at increasing effector-to-target (E:T) ratios. αCDH3-CAR T cells demonstrate limited lysis of CDH3-knockout cells, confirming CDH3 dependency cytotoxicity.

### Figure S7. CDH3 targeted CAR T in vivo

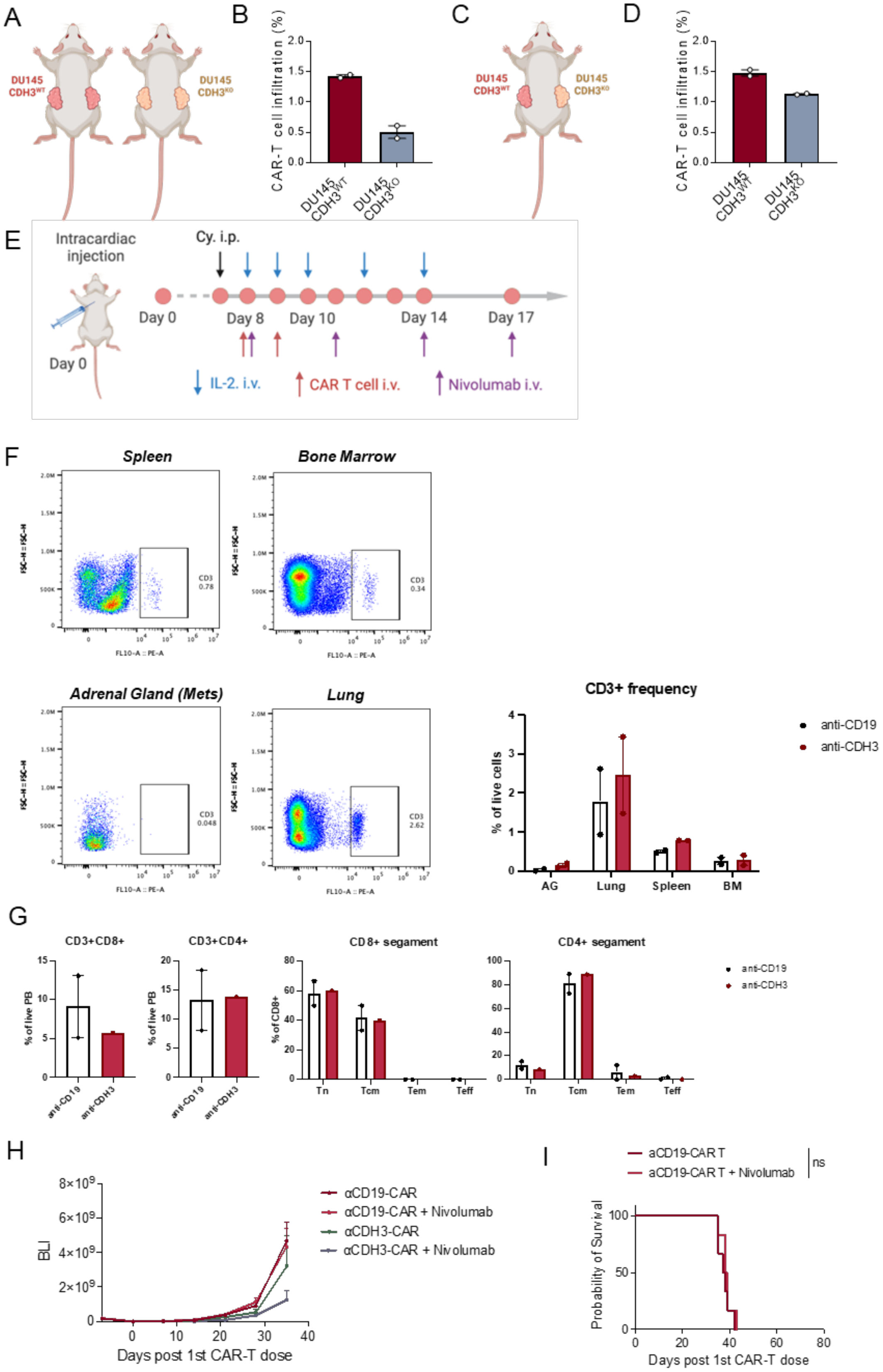

(A) NOD/SCID mice were inoculated subcutaneously with DU145-CDH3$^{WT}$ or DU145-CDH3$^{KO}$.

(B) After 3 weeks of tumor growth, αCDH3-CAR T cells were administered intravenously (with Cy. precondition and IL-2 support), and tumors were harvested 7 days later to exam the CAR T cell infiltration.

(C) NOD/SCID mice were inoculated subcutaneously with DU145-CDH3$^{WT}$ on one flank and DU145-CDH3$^{KO}$ on the contralateral flank.

(D) After 3 weeks of tumor growth, αCDH3-CAR T cells were administered intravenously (with Cy. precondition and IL-2 support), and tumors were harvested 7 days later to exam the CAR T cell infiltration.

(E) Schematic of the in vivo experimental design metastatic prostate cancer models for the two rounds of the CAR-T therapy. DU145-TR are delivered via intracardiac injection on Day 0, followed by cyclophosphamide (Cy) administered intraperitoneally (i.p.) and supportive treatments (IL-2), along with separate or combined infusions of CAR T cells and nivolumab.

(F) From the same batch of mice, different tissue (adrenal glands with metastasis) were harvested to test the infiltration level

(G) NOD/SCID mice were intracardiac injected with DU145-TR to build the similar metastatic prostate cancer model. After 4 weeks of tumor growth, αCD19-CAR T and αCDH3-CAR T cells were administered intravenously (with Cy. precondition and IL-2 support), and mice were sacrificed 1 days later to exam the proportion of CD8 vs. CD4 population and corresponding T-cell subsets (naïve, T effector, T effector memory, and T central memory) in the peripheral blood.

(H) Corresponding to the Figure 7D. BLI signals of the DU145-TR metastatic prostate cancer mode, comparing αCD19-CAR T alone, αCD19-CAR T + nivolumab, αCDH3-CAR T alone, and αCDH3-CAR T + nivolumab over time.

(I) Survival curves of αCDH19-CAR T alone and αCDH19-CAR T + nivolumab treatment groups.
